# Molecular competition can shape enhancer activity in the Drosophila embryo

**DOI:** 10.1101/2021.05.07.443186

**Authors:** Rachel Waymack, Mario Gad, Zeba Wunderlich

## Abstract

Transgenic reporters allow the measurement of regulatory DNA activity *in vivo* and consequently have long been useful tools in the study of enhancers. Despite the utility of transgenic reporters, few studies have investigated the potential effects these reporters have on the expression of other transgenic reporters or endogenous genes. A full understanding of the impacts transgenic reporters have on expression is required for accurate interpretation of transgenic reporter data and characterization of gene regulatory mechanisms. Here, we investigate the impact transgenic reporters have on the expression of other transgenic reporters and endogenous genes. By measuring the expression of *Kruppel* (*Kr*) enhancer reporters in live *Drosophila* embryos that contain either one or two copies of identical reporters, we find reporters have an inhibitory effect on one another’s expression. Further, expression of a nearby endogenous gene is decreased in the presence of a *Kr* enhancer reporter. Through the use of competitor binding site arrays, we present evidence that reporters, and potentially endogenous genes, are competing for transcription factors (TFs). Increasing the number of competitor Bcd binding sites decreases the peak levels and spatial extent of Bcd-regulated enhancer reporters’ expression. To understand how small numbers of added TF binding sites could impact gene expression to the extent we observe, we develop a simple thermodynamic model of our system. Our model predicts competition of the measured magnitude specifically if TF binding is restricted to distinct nuclear subregions, underlining the importance of the non-homogenous nature of the nucleus in regulating gene expression.

## Introduction

An organism’s ability to precisely control gene expression is dependent on the activity of enhancers. Through the binding of specific combinations of transcription factors (TFs), which can be activating or repressive, enhancers are able to control the expression of their target genes in time and space. Enhancers control gene expression across all aspects of organismal functioning, from the immune system to the nervous system, and play a particularly important and well-studied role in the process of embryonic development (Levine, 2010; Shlyueva, et al., 2014). During this period, enhancers regulate the expression of genes that determine critical cell fate decisions underlying patterning and organogenesis.

A significant amount of our understanding of enhancers and other *cis*-regulatory elements has come from the use of transgenic reporter lines. These transgenic animals have measurable reporters, such as fluorescent proteins or LacZ, under the control of *cis*-regulatory elements to enable observation of that element’s activity in living organisms in different life stages, tissue types, or conditions (reviewed in Kvon, 2015; Wood, 1995). Studies of transgenic animals have enabled the discoveries of previously unknown enhancers, the modularity of enhancers, and the importance of the arrangement of TF binding sites or enhancer “grammar”, among others (Pennacchio, et al., 2006; O’Kane & Gehring, 1987; Bier, et al., 1989; Visel, et al., 2009; Swanson, et al., 2010).

Despite the remarkable utility of transgenic reporters, or perhaps in part because of it, little work has been done to look at the effect of these reporters on expression of other reporters or endogenous genes. Although reporters are exogenous regions of DNA that can originate from completely different species than the host animal, once integrated into the genome, these transgenes rely on the same pool of transcription factors, polymerases, and other molecular factors required for gene expression as endogenous genes. Given that most of these factors are present at relatively high copy numbers in the cell, for example 250,000 Zld TF molecules per nucleus in the *Drosophila* embryo (Biggin, 2011; BNID 106849, Milo et al., 2010) or over 80,000 RNA polymerase (RNAP) molecules per nucleus in human cells (Zhao, et al., 2014; BNID 112321, Milo et al., 2010), it is commonly assumed that adding an additional enhancer would have little impact on the availability of key expression machinery. However, a couple of examples suggest that there may be competition between transgenic reporters and endogenous genes. A study by Laboulaye, et al. measured the effect of three different transgenic reporters on endogenous gene expression in mice (Laboulaye, et al., 2018). The authors found that the transgenic reporters all decreased the expression of the closest endogenous gene. Thompson & Gasson noted that endogenous protein levels may be slightly decreased in *Saccharomyces cerevisiae* and *Lactococcus lactis* expressing transgenic reporters, but the results were inconclusive (Thomspon & Gasson, 2001). These examples suggest that transgenic reporters may decrease endogenous gene expression but leave open the questions of the mechanisms behind these decreases and whether such an effect is limited to certain organisms or reporters.

Like much of the field, we often used transgenic reporters under the assumption that they had no effect on the expression of other genes until we saw evidence to the contrary in our own data. In a study investigating gene expression noise in *Drosophila* embryos, we observed evidence of competition between identical copies of transcriptional reporters integrated on homologous chromosomes (Waymack, et al., 2020). We were surprised to find that homozygous reporter embryos produced less mRNA per reporter allele than hemizygous embryos, with a reporter present on only one of the two homologous chromosomes (Figure 1). We suspected this could have important implications not only for the use of transgenic reporters, but also for our understanding of the balance between the supply of and demand for transcriptional machinery within the nucleus.

**Figure 1 –.**
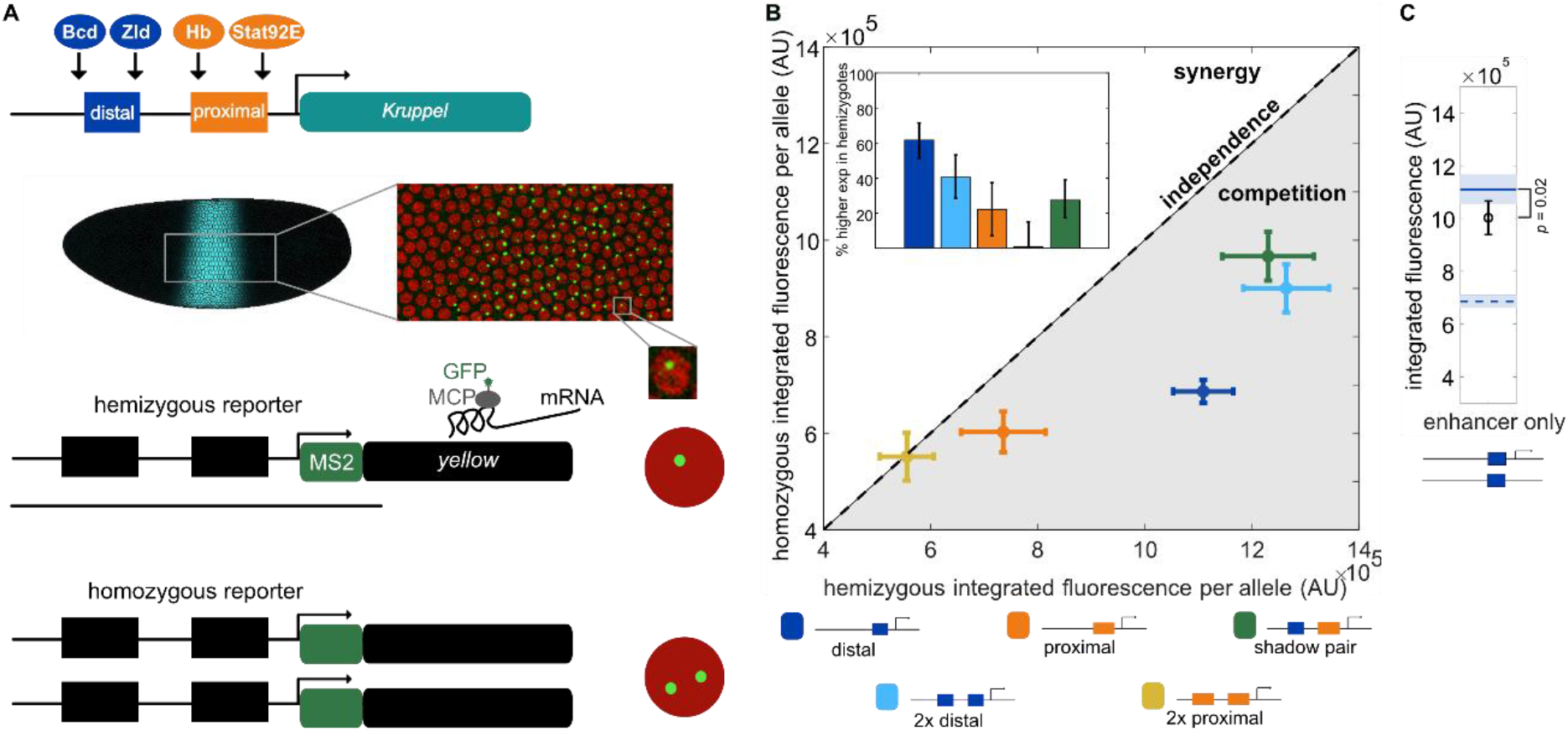
Differences in mRNA production in homozygous and hemizygous embryos suggest competition between reporters. **A**. This panel is adapted from Waymack, et al. (Waymack, et al., 2020). (Top) *Kr* expression in the early embryo is controlled by the activity of a pair of shadow enhancers, termed proximal and distal based on their location relative to the *Kr* promoter, that are each activated by different transcription factors (TFs). (Middle) The expression pattern driven by this pair of shadow enhancers is a stripe in the center 20% of the embryo. We use the MS2 system to image active transcription driven by enhancer reporters in living embryos. The cut out from embryo shows a still frame of a movie where red circles are nuclei and green spots are sites of active transcription. To test whether transgenic reporters affect each other’s expression, we generated embryos that are either homozygous or hemizygous for a particular reporter construct. (Bottom) Hemizygous embryos have the enhancer-MS2 reporter inserted on only one homologous chromosome and therefore display one transcriptional spot per nucleus. Homozygous embryos have the same enhancer-MS2 reporter inserted at the same location on both homologous chromosomes and therefore display two transcriptional spots per nucleus. **B**. mRNA production from homozygous reporter constructs compared to production from hemizygous constructs suggests competition between reporters. The graph shows total mRNA produced per allele in homozygous embryos as a function of total mRNA produced per allele in hemizygous embryos for the reporter construct indicated. The dashed diagonal line represents expected expression assuming independent activity of the two reporters in homozygous embryos. Points falling above this line display synergy in the activity of the two reporters in homozygotes and points falling below display competition. Error bars indicate 95% confidence intervals. Inset shows the percent higher expression in hemizygous versus homozygous embryos for each reporter construct. Error bars represent 95% confidence intervals from 1000 rounds of bootstrapping. **C**. To rule out reporter competition being an artifact of our imaging system, we measured expression of the distal enhancer MS2 reporter in the presence of a non-transcribing distal enhancer on the homologous chromosome. The second distal enhancer is identical to the reporter, but lacks both a promoter and MS2 sequence. The graph shows expression driven by the distal enhancer reporter is significantly reduced in the presence of the non-transcribing distal enhancer (*p*-value = 0.02, t-test). The top (solid) horizontal line indicates expression driven by the distal enhancer reporter in the hemizygous configuration and the bottom (dashed) horizontal line indicates peak expression per allele driven in the homozygous configuration. Error bars and shading represent 95% confidence intervals.

Here we track the activity of multiple configurations of transgenic reporters in *Drosophila* embryos to assess the impact of these reporters on one another and endogenous genes. We measured live mRNA dynamics driven by the embryonic enhancers of the gap gene *Kruppel* (*Kr*) in the presence or absence of a second transcriptional reporter or a competitor TF binding array. We find that enhancer reporter expression is lower not only in the presence of a second reporter, but also in the presence of non-transcribing TF binding arrays, suggesting that there is competition for locally limited levels of certain TFs. This effect is not restricted to reporters; expression of a nearby endogenous gene is also decreased in transgenic embryos. To understand how the addition of the relatively small number of TF binding sites present in our constructs can measurably decrease reporter expression, we developed a thermodynamic model of our system. We predict reduced expression of the magnitude observed in transgenic embryos if we assume TF binding is restricted to so-called “hub” regions, but not if we assume TFs have access to the whole genome. This work reconciles the question of how tens of TF binding sites in a transgenic reporter construct can impact the available supply of tens of thousands of TF molecules. We suggest that the TF supply relevant to a particular enhancer is limited to a smaller pool of the TFs in a nucleus.

## Results

### Homozygous reporters display evidence of competition

To test whether transgenic reporters affect the expression of other alleles, we compared the expression output in embryos either homozygous or hemizygous for different reporter constructs. In the absence of reporter interactions, we expect to see the same levels of mRNA production per allele in hemizygous embryos and homozygous embryos. Conversely, if the reporters do affect one another’s expression, then expression levels per allele will differ in hemizygous vs homozygous embryos, depending on the nature of this interaction. A synergistic interaction, perhaps through a mechanism such as increasing the local concentration of a key TF, would lead to higher levels of transcription in homozygous embryos than hemizygous embryos (Figure 1B upper half). An antagonistic interaction, such as competition for a limited shared resource, would lead to lower levels of transcription in homozygous embryos than hemizygous embryos (Figure 1B lower half).

To assess the nature of potential reporter interactions, we measured transcriptional output of different enhancers in living embryos using the MS2 reporter system. When transcribed, the MS2 sequence forms stem loops that are then bound by an MCP-GFP fusion protein expressed in the embryo, enabling us to visualize sites of nascent transcription (Figure 1A; Garcia, et al., 2013). We can track these individual transcriptional spots across the time of nuclear cycle 14 (nc14), when these enhancers are most active, to measure total transcriptional output and dynamics. As a test case, we used different combinations of the two *Kruppel* (*Kr*) embryonic shadow enhancers. The *Kr* shadow enhancer pair, together or individually, drives a stripe of expression in the central 20% of the embryo (Figure 1A). We generated transgenic flies with each individual enhancer, the shadow enhancer pair, or each enhancer duplicated in tandem driving an MS2 reporter (Figure 1B). Despite the similar pattern of expression driven by the two individual enhancers, the distal and proximal enhancers are each activated by different sets of TFs (Wunderlich, et al., 2015). We previously showed that this separation of TF inputs plays an important role in suppressing gene expression noise (Waymack, et al., 2020). Here, this separation of TF inputs allows us to investigate whether the reporter interactions we observe are influenced by specific regulatory factors or are more general consequences of having two reporters present.

In the majority of cases, hemizygous embryos produce more mRNA per allele than do homozygous embryos (Figure 1B). To calculate the mRNA produced by each reporter, we integrate the area under the fluorescence traces of activity measured during nc14 at the anterior-posterior position in the embryo of peak expression (Supplemental Figure 1). The single and duplicated distal constructs produce 62% and 40%, respectively, more mRNA per allele in hemizygous embryos than in homozygous embryos. The shadow pair and proximal enhancer reporters produce 27% and 22% more mRNA per allele at their respective regions of peak expression in hemizygous embryos than in homozygous embryos. The duplicated proximal construct drives the same level of expression in hemizygous and homozygous embryos. By comparing the competition exhibited by duplicated and single enhancers, we do not find evidence that longer reporter sequences drive stronger reporter competition (Figure 1B inset).

We suspect this trend may arise because duplicated enhancers with a large array of similar binding sites can recruit a larger pool of TFs (Tsai, et al., 2019) or because there can be synergy between the enhancers in promoter activation (Bothma, et al., 2015). In sum, when two reporters are present in the same nucleus, neither typically transcribes to its full potential, suggesting that there is some form of competition between the two reporters. We hypothesized that the reporters are competing for one or more molecular factors required for reporter transcription or visualization.

### Reporter competition is not an artifact of imaging system

To assess whether reporter competition is the result of a biological phenomena, such as limiting levels of a TF, or an artifact of our reporter system, such as limiting levels of MCP-GFP, we measured reporter output in the presence of a second non-transcribing transgenic construct. We produced a version of our distal enhancer construct that lacks both a promoter and the MS2 cassette. This construct therefore can bind the same regulatory TFs as the original distal construct but will not drive transcription. Therefore, it should not interact with promoter-bound factors, such as RNAP, or the MCP-GFP coat protein. If the observed competition is for MCP-GFP or is dependent on transcription, we expect to see no effect on reporter expression when the enhancer-only construct is present on the homologous chromosome. Conversely, if one or more regulatory factors binding the enhancer is limiting, we expect to see a decrease in reporter expression, similar to the lower expression of homozygous versus hemizygous embryos (Figure 1B).

In the presence of the distal enhancer-only construct, the distal enhancer reporter drives 11% lower levels of expression at its region of peak expression than in the hemizygous configuration (Figure 1C). While significant (t-test *p*-value = 0.02), this decrease is not as large as the one we see when a second transcribing distal enhancer reporter is present on the homologous chromosome. We suspect the smaller effect of the non-transcribing distal enhancer construct is due to differences in the exact composition and levels of factors that are recruited to transcriptionally active versus inactive enhancers (Savic, et al, 2015; Bozek & Gompel, 2020; Li, et al., 2008). To further rule out that the observed reporter competition is a result of limiting levels of the MCP-GFP reporter, we looked at the pattern of reporter competition across the length of the embryo. The MCP-GFP coat protein is expressed ubiquitously across the length of the embryo, while many of the TFs regulating the *Kr* enhancers are spatially patterned. If reporters are competing for limited levels of MCP-GFP, we would expect to see the highest rates of competition in the center of the embryo where expression driven by our reporters is highest (Supplemental Figure 2B). Instead, with all of our reporter constructs we find that rates of competition are highest outside of the region of peak expression (Supplemental Figure 2C-E), strongly suggesting that reporters are not competing for limited levels of MCP-GFP. Instead, this pattern of competition rates combined with the finding that a non-transcribing enhancer construct can reduce reporter activity suggest that our reporters are competing for an endogenous factor required for transcription.

### Reporters are competing for transcription factors

While reporter competition seems to be independent of the MS2 system or weakly dependent on transcription itself, it does depend on the identity of the enhancer driving reporter expression (Figure 1B). The presence of a second identical reporter has a large effect on expression driven by the shadow construct and an even larger effect on the duplicated distal construct, while it has no significant effect on the duplicated proximal construct (Figure 1B). Since the *Kr* distal and proximal enhancers are regulated by separate sets of TFs (Wunderlich, et al., 2015), we hypothesized that reporters may be competing for one or more of these TFs and that this may underlie the difference in competition levels between constructs. To test this hypothesis, we measured the effect of TF binding site arrays on the activity of the reporters. As the level of competition is not significantly different between the single and duplicated enhancer constructs, we focused on the two duplicated enhancers and the shadow pair, which are similar lengths and therefore have similar numbers of TF binding sites. We created DNA sequences consisting of six strong TF binding sites for each of the key activating TFs of the *Kr* enhancers and inserted them into the identical site on the homologous chromosome opposite one of the enhancer-MS2 reporters (Figure 2A). Critically, these TF binding site arrays lack promoter and MS2 sequences. We reasoned that these TF binding site arrays would function to sequester TF molecules without affecting factors specifically involved in transcript production (such as RNAP) or reporter visualization (i.e. MCP-GFP). Therefore, any changes in transcriptional output by the enhancer-MS2 reporter observed in the presence of a TF binding site array should stem from decreased levels of available TF, not higher demand for basal transcriptional machinery or MCP-GFP.

**Figure 2 –.**
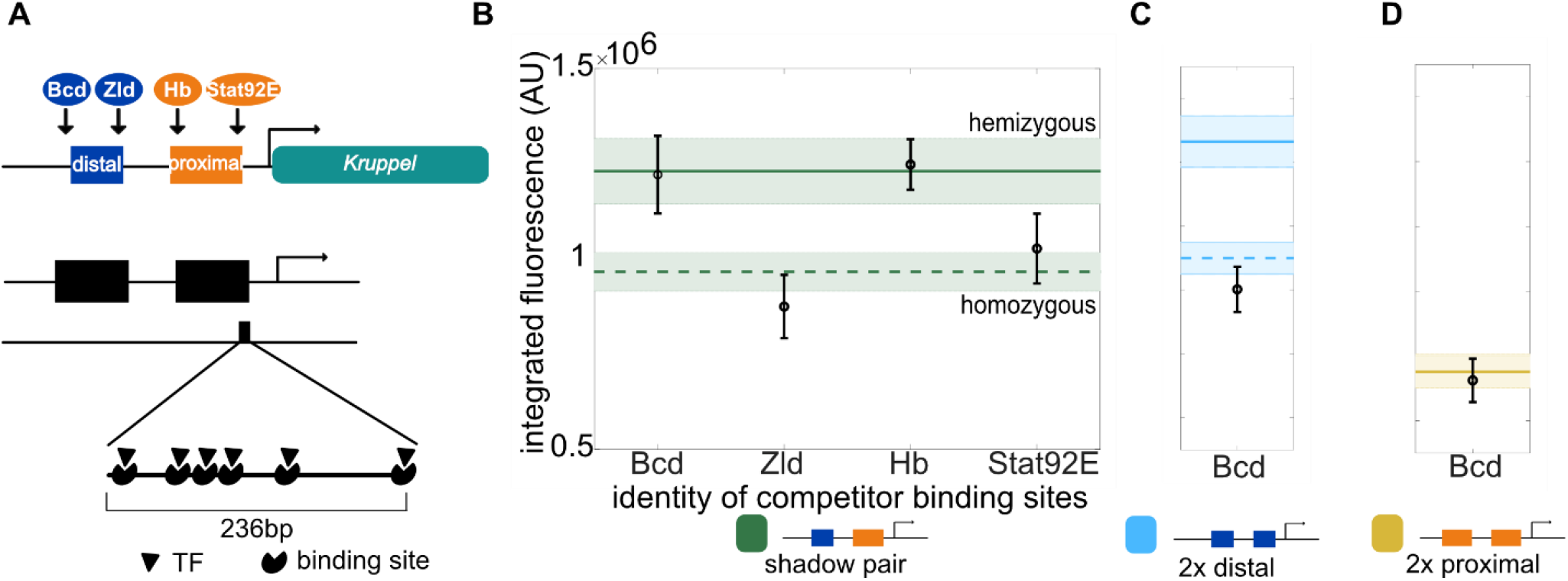
Competitor TF binding sites on homologous chromosome decrease reporter activity. To test whether limiting levels of one or more activating TFs contribute to the reporter competition we observe, we measured the activity of our reporters in the presence of TF binding site arrays. **A**. (Top) The *Kr* shadow enhancers are activated by different sets of TFs. (Bottom) A schematic of TF binding site arrays that are intended to act as sinks for TF molecules. The arrays are each 236bp long, contain six binding sites for the indicated TF, and are inserted at the same genomic site as enhancer-MS2 reporters on the homologous chromosome. The binding site arrays do not contain a promoter or MS2 sequence. **B**. The activity of the shadow pair reporter is reduced in the presence of some TF binding site arrays. Graph shows the peak expression of the shadow pair in the presence of the indicated TF binding site array on the homologous chromosome. In B-D, the horizontal solid line indicates the peak expression level in hemizygous embryos of the indicated reporter construct and the horizontal dashed line indicates the peak expression level per allele in homozygous embryos. **C**. The activity of the duplicated distal reporter is reduced to homozygous levels when the Bcd binding array is present on the homologous chromosome. **D**. Activity of the duplicated proximal reporter, which is not activated by Bcd, is not reduced when the Bcd binding array is present on the homologous chromosome. Note that the homozygous and hemizygous peak expression levels (dashed and solid horizontal lines) overlap for the duplicated proximal reporter. Error bars and shading in B-D indicate 95% confidence intervals.

Specifically, we created four binding site arrays corresponding to the four key TF activators of the shadow pair (Bicoid (Bcd), Hunchback (Hb), Stat92E, and Zelda (Zld); Figure 2A), which each contain six binding sites for the respective TF across 236bp. As the shadow pair is the only construct known to be regulated by all four TFs, we first assessed the impact of these binding site arrays on the activity of the shadow pair reporter. We find that the binding site arrays for the Zld and Stat92E each reduce the activity of the shadow pair down to the levels seen in hemizygous embryos, while the Bcd and Hb binding site arrays do not have a significant effect on the shadow pair’s activity (Figure 2B). We suspect that the Stat92E and Zld binding arrays may have the largest effect on the shadow pair’s activity due to their essential roles in early gene activation (Harrison, et al., 2011; Tsurumi, et al., 2011).

As we observe a stark difference in the levels of competition in the duplicated distal versus duplicated proximal constructs, we asked whether the Bcd binding site array affects expression of either construct. Bcd is a key activator of the distal enhancer, but not the proximal enhancer (Figure 2A). In line with this, the duplicated distal reporter’s activity is reduced 37% compared to hemizygous levels at their regions of peak activity in the presence of the Bcd binding site array (Figure 2C), while the activity of the duplicated proximal enhancer is not significantly changed (Figure 2D). The large effect the Bcd array (from here on called 1xBcd) on the duplicated distal reporter is striking, as the TF binding site array is less than one-fifth the size of either *Kr* enhancer and contains only six, albeit strong, binding sites for Bcd. The specificity of the 1xBcd array in reducing expression only of the Bcd-activated duplicated distal reporter, but not of the duplicated proximal reporter, suggests that the effect we observe is specific to sequestering Bcd molecules, and not a general effect of inserted DNA sequences.

### Reporters show dosage-dependent response to increasing number of Bcd competitor sites

As a whole, these experiments suggest that limiting levels of TFs play an important role in reporter competition. When comparing the effects of the 1xBcd array across constructs, the expression of the duplicated distal reporter is dramatically reduced in the presence of the array, while the expression of the duplicated proximal and shadow pair constructs are unaffected.

Given that the shadow pair is regulated by more TF inputs beyond Bcd than is the duplicated distal reporter, we hypothesized that the shadow pair may be less sensitive to Bcd competition. To test this hypothesis, we attempted to sequester larger amounts of Bcd and measure the effect on the shadow pair’s activity. We measured the activity of the shadow pair reporter in the presence of larger binding site arrays consisting of three (3xBcd; 3 x 6 copies = 18 Bcd binding sites) or six (6xBcd; 6 x 6 = 36 Bcd binding sites) copies of the original Bcd binding site array.

In line with our hypothesis, we find that the shadow pair reporter activity decreases with increasing number of Bcd binding sites in the competitor array. Peak expression is reduced 1% in the presence of the 1xBcd array, 21% with the 3xBcd array, and 38% with the 6xBcd array relative to expression in hemizygotes (Figure 3A). We also measured expression of the duplicated distal enhancer with the larger Bcd binding arrays and find a non-linear effect of increasing the number of competitor Bcd binding sites (Supplemental Figure 3A; Discussion).

**Figure 3 –.**
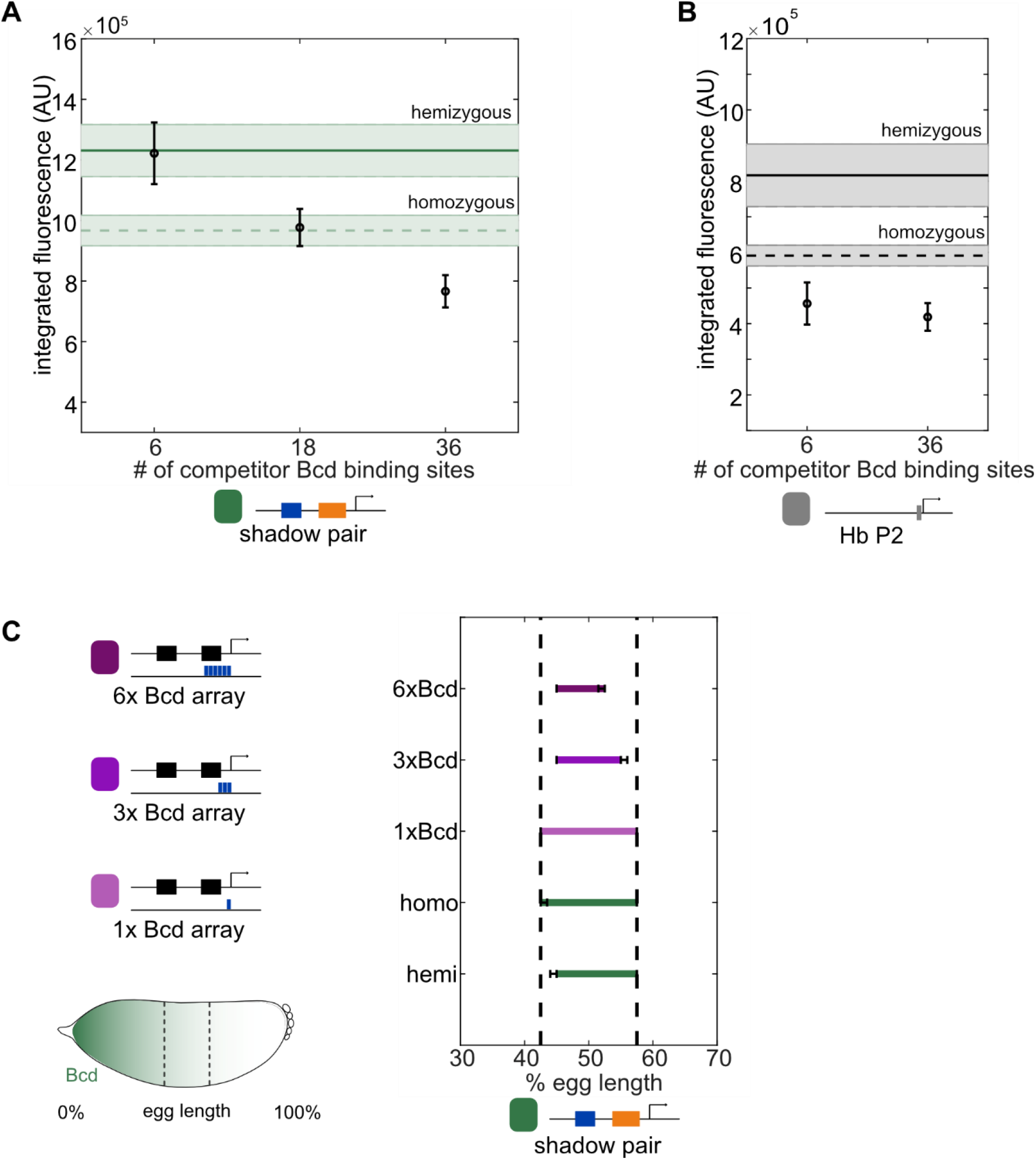
Competitor Bicoid binding sites decrease and shift the activity of shadow pair reporter. To assess whether limiting levels of the activating TF Bicoid (Bcd) cause the apparent competition between reporters observed, we measured the transcriptional output of the shadow pair construct in the presence of Bcd binding site arrays of increasing length on the homologous chromosome. The 1xBcd binding site array consists of six Bcd binding sites but lacks a promoter or MS2 cassette. The 3xBcd and 6xBcd binding site arrays are three and six repeats, respectively, of the 1xBcd array and therefore contain a total of 18 and 36 Bcd sites, while also both lacking a promoter or MS2 cassette. These binding site arrays were inserted into the same location on Chromosome 2 as the enhancer reporters. **A**. Peak expression per allele driven by the shadow pair reporter decreases as the number of competitor Bcd binding sites increases. The horizontal lines mark the peak total expression per allele driven by the shadow pair reporter as hemizygotes (top solid line) or homozygotes (bottom dashed line). Shading and error bars indicate 95% confidence intervals. **B**. Competitor Bcd binding site arrays decrease the expression of an unrelated Bcd-responsive enhancer. To test if the effect of the Bcd binding arrays is specific to the *Kr* enhancers, we measured expression driven by the *hunchback* P2 (*hb*P2) enhancer, which is also activated by Bcd, in the presence of the 1x and 6xBcd binding site arrays. The graph shows the peak expression driven by the *hb*P2 reporter in the presence of the indicated Bcd binding site arrays. Shading and error bars in A and B indicate 95% confidence intervals. **C**. (Left) Bcd is expressed in a gradient from the anterior of the embryo (0% egg length) to the posterior (100% egg length). The *Kr* expression domain is indicated by dashed vertical lines. Schematics above the embryo diagram show the 1x, 3x, or 6xBcd arrays used with the enhancer reporters. (Right) Expression patterns driven by the shadow pair reporter in the presence of increasing numbers of competitor Bcd binding sites. Graph shows the range of the expression pattern of each configuration to 50% of peak expression levels of the homozygous configuration, whose boundaries are indicated with dashed vertical lines. Error bars represent 95% confidence intervals found from 1000 rounds of bootstrapping.

To assess whether the reduction in mRNA output in the presence of the Bcd array is specific to the *Kr* enhancers or a general phenomenon, we measured the expression driven by the *hunchback* (*hb*) P2 Bcd-responsive enhancer in the presence or absence of the 1xBcd and 6xBcd arrays. Similar to our findings with the *Kr* enhancers, the Bcd binding arrays decrease the expression of the *hb* P2 enhancer (Figure 3B). Relative to hemizygous levels, peak expression of the *hb* P2 enhancer is decreased 44% with the 1xBcd array and 49% with the 6xBcd array. We suspect that the relatively modest effect of larger numbers of Bcd competitor sites on reporter activity stems from an upper limit to the amount of Bcd molecules that can be effectively sequestered away from the enhancers at our binding site arrays along with the activating function of other TFs.

Since TFs can control both the level and pattern of enhancer activity, we measured how the expression boundaries of our reporters changed in response to the Bcd binding arrays. Bcd is expressed in a gradient from the anterior to the posterior of the embryo. Even though Bcd is present at high levels in the anterior of the embryo, *Kr* enhancers do not drive expression there because of repression by Giant (Gt), Knirps (Kni), and Hb, which can act as both an activator and a repressor (Jaeger, 2011; Vincent, et al., 2018; Papatsenko & Levine, 2008). Therefore, we expected little effect on the anterior boundary by the Bcd binding site arrays. In contrast, since the posterior boundary is partially set by the decreasing levels of Bcd, if our Bcd arrays functionally reduce Bcd levels available for enhancer activation, we would expect to see a larger effect at the posterior boundary. We find that the posterior boundary of the shadow pair’s expression domain moves towards the anterior in response to increasing number of competitor Bcd binding sites (Figure 3C). Relative to the homozygous configuration, the posterior border of shadow pair expression shifts anteriorly 2.5% of embryo length in the presence of the 3xBcd array and 5% in the presence of the 6xBcd array. Similar to peak expression levels, the 1xBcd array does not change the expression boundaries of the shadow pair reporter. We see similar anterior shifts of the duplicated distal expression pattern with the Bcd binding arrays that qualitatively match the decrease in peak expression seen with each array (Supplemental Figure 3B). We note that the anterior boundary shifts towards the posterior in the presence of the 3xBcd and 6x Bcd arrays, which we suspect stems from the balance of activity between Bcd and the repressive TFs in this region (Kraut & Levine, 1991; Papatsenko & Levine, 2008; Small, et al., 1991; Stanojevic, et al., 1991).

### Competition occurs at another genomic site and with an endogenous gene

Based on our findings thus far suggesting that reporter competition stems from competition for Bcd and other TFs, we reasoned that this competition should occur at other genomic insertion sites and with endogenous genes reliant on the same TFs. To first assess whether the observed reporter competition occurs at other genomic insertion sites, we measured the expression of the reporters in homozygous versus hemizygous configurations when inserted into a different chromosome (chromosome 3). Similar to our findings at the chromosome 2 insertion site (Figure 1), expression levels driven by the duplicated distal and shadow pair reporters are significantly lower in the presence of a second identical reporter (Figure 4A, B). On chromosome 3, expression in homozygous embryos is 82% and 75% of expression in hemizygous embryos for the duplicated distal and shadow pair reporters, respectively. With both of these reporters, the degree of competition is consistent between the two insertion sites, indicating that the observed competition occurs at different genomic locations (Figure 4 A and B insets).

**Figure 4 –.**
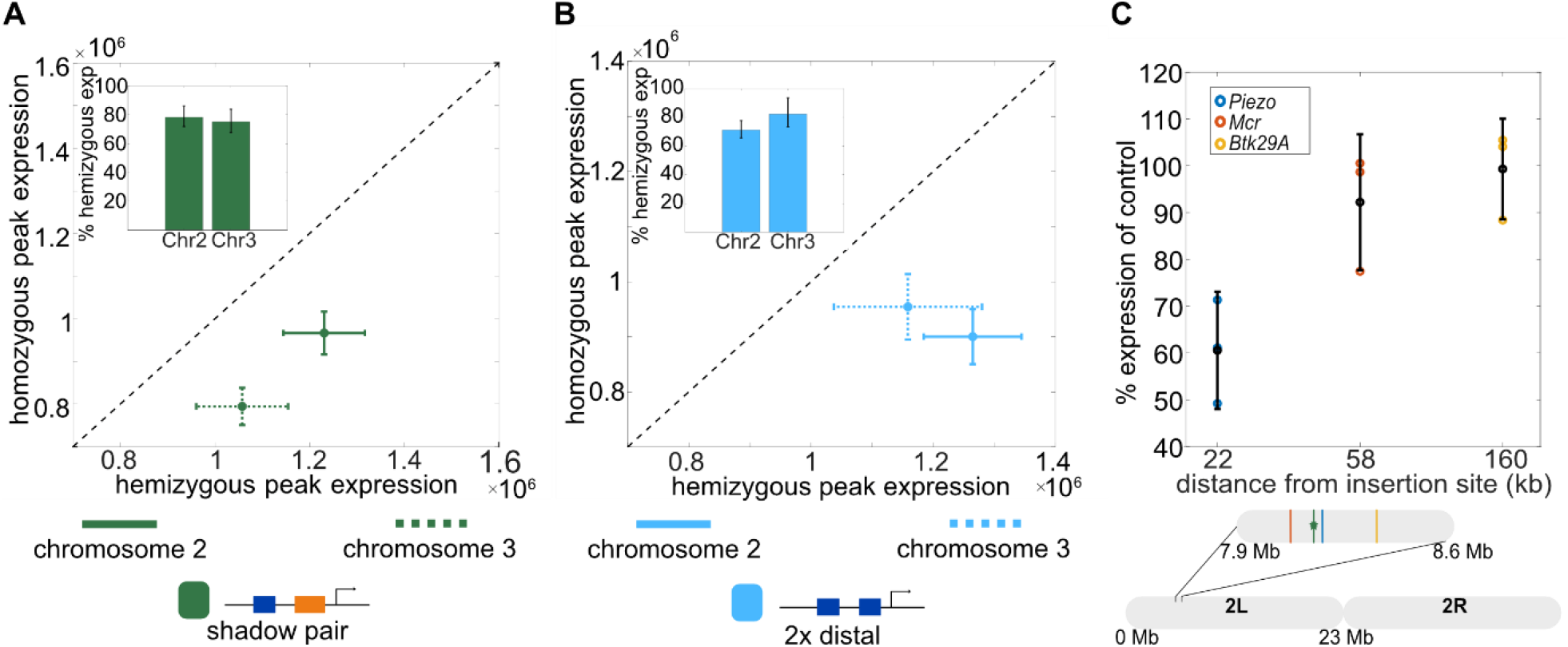
Competition occurs at an additional location and gene. Based on our data suggesting that reporters are competing for limited levels of TFs, we suspected this competition would also occur at other transgenic insertion sites and with endogenous genes. **A**. Reporter competition occurs at multiple genomic insertion sites. Graph shows the peak expression levels per allele in homozygous embryos as a function of the peak expression levels in hemizygous embryos for the shadow pair reporter inserted in either chromosome 2L or 3L. The data for chromosome 2 are the same as in Figure 1B. Diagonal line marks expected values for homozygous expression if reporters do not interact and instead display independent expression. Error bars represent 95% confidence intervals. Inset shows the peak expression levels in homozygous embryos relative to hemizygous embryos with the shadow pair reporter inserted on either chromosome 2 or chromosome 3. Error bars in inset represent 95% confidence intervals from 1000 rounds of bootstrapping. **B**. The graph is as in A with the duplicated distal reporter inserted on chromosome 2 or chromosome 3. **C**. To determine the effect, if any, of transgene’s use of resources on endogenous genes’ expression, we compared the expression levels of three endogenous genes likely to be Bcd-regulated at increasing genetic distances from the transgenic insertion site in embryos with or without the duplicated distal transgene. Graph shows the fold change in expression of *Piezo, Mcr*, and *Bkt29A* in embryos homozygous for the duplicated distal transgene compared to WT embryos as measured by qPCR. Error bars represent 95% confidence intervals and black circles indicates the mean. Schematic below graph shows the genetic distance of the three measured genes (indicated with a blue, red, or yellow vertical line) from the attP site (VK000002) on chromosome 2L (marked with green line and star) where all transgenic constructs, unless otherwise specified, were inserted.

Based on previous work in the mouse, we suspected that the reporter-induced competition would be limited to endogenous genes that are a short linear distance from the reporter insertion site (Laboulaye, et al., 2018). To assess whether this is true, we measured the expression of three genes likely to be regulated by Bcd at varying linear distances from the chromosome 2 insertion site. We measured the expression of *Piezo* (22kb from insertion site), *Mcr* (58kb from insertion site), and *Btk29A* (160kb from insertion site) via qPCR in embryos with or without two copies of the duplicated distal transgene. All three of these genes are predicted to be regulated by Bcd, based on both previously measured expression patterns and Bcd binding near these genes in the early embryo (Fisher, et al., 2012; Hannon, et al., 2017; Supplemental Figure 4). In transgenic embryos, expression of the gene closest to the insertion site, *Piezo*, is significantly reduced to 60% of the levels seen in embryos of the same genetic background but lacking the transgene (Figure 4B). The expression levels of *Mcr* and *Btk29A*, which are further removed from the transgenic insertion site, are not significantly changed in transgenic embryos (Figure 4B). This potential distance-dependent effect of our transgene on endogenous gene expression is consistent with our finding that, in homozygous reporter embryos, there is more competition in nuclei in which the MS2 spots are physically closer together (Supplemental Figure 5B).

### A hub-based model of TF-enhancer interactions predicts TF competition

We were initially surprised to find that a reporter construct, with a length less than 0.001% of the genome, can have measurable effects on the expression of both other reporters and a nearby endogenous gene. Even more surprising is our finding that Bcd binding site arrays, which do not themselves drive any expression and have as few as six binding sites, also significantly reduce the expression of our Bcd-regulated enhancer reporters (Figure 2 & 3). This suggests that competition for Bcd can be induced by the addition of a relatively small number of binding sites, despite the fact that Bcd copy numbers vary between approximately 1500 and 3000 molecules per nucleus in the region of *Kr* expression (Biggin, 2011; Fowlkes, et al., 2008). To better understand how the addition of a small sequence could induce competition for TFs, we developed a simple thermodynamic model of our system. The goal of our modeling effort is not to fit parameters such that the model precisely recapitulates our experimental data, but rather to see if our experimental observations are sensible by generating ballpark estimates of molecular competition using models that only rely on measured biophysical parameters.

Our model predicts the probability of a TF being bound to a target site, such as one of the binding sites that exist in an enhancer (Figure 5). For simplicity, we assume that TF binding at the target site is proportional to enhancer activity (Bintu, et al., 2005; Phillips, et al., 2019). In reality, enhancer activity depends on the combined occupancy of many TF binding sites (Levine, 2010). The simplifying assumption that enhancer activity is proportional to binding site occupancy allows us to avoid the need to test multiple models with different components, such as cooperative TF binding or activation behavior. In addition to the target site, TF molecules can bind to specific or non-specific competitor sites. Since most TFs have sequence-independent affinity for DNA (Slattery, et al., 2014), the number of non-specific binding sites, *N*, is set to 1×10^8^, roughly the size of the *Drosophila melanogaster* genome. The number of specific competitor sites, *C*, is varied. To maintain the simplicity of the model, the binding energy of specific competitor sites, *E*_*s*_, is equal to the binding energy of the target site, while the binding energy of all non-specific sites is represented as *E*_*ns*_. Since the specific binding energy is representing that of multiple binding sites, which may differ in their affinities, we vary the difference between specific and non-specific binding energies. Lastly, to allow for comparisons with our experimental data and to measure the effect of TF levels on binding, we vary the levels of our input TF *T* as a function of embryo length, *l*, in accordance with the measured Bcd gradient (Fowlkes, et al., 2008). In this way, we can look at how the probability of TF binding to a single target site, *p*(*bound; T(l)*), changes as a function of number of specific competitor sites, binding strength relative to non-specific binding, and TF abundance.

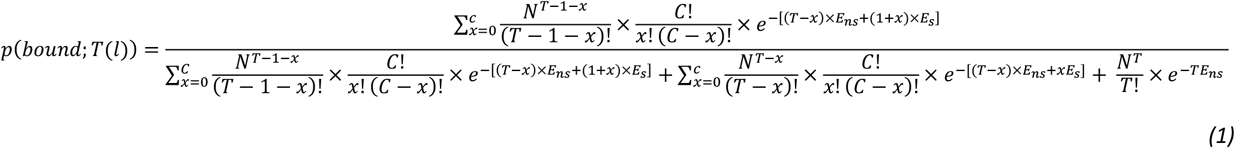

**Figure 5 –.**
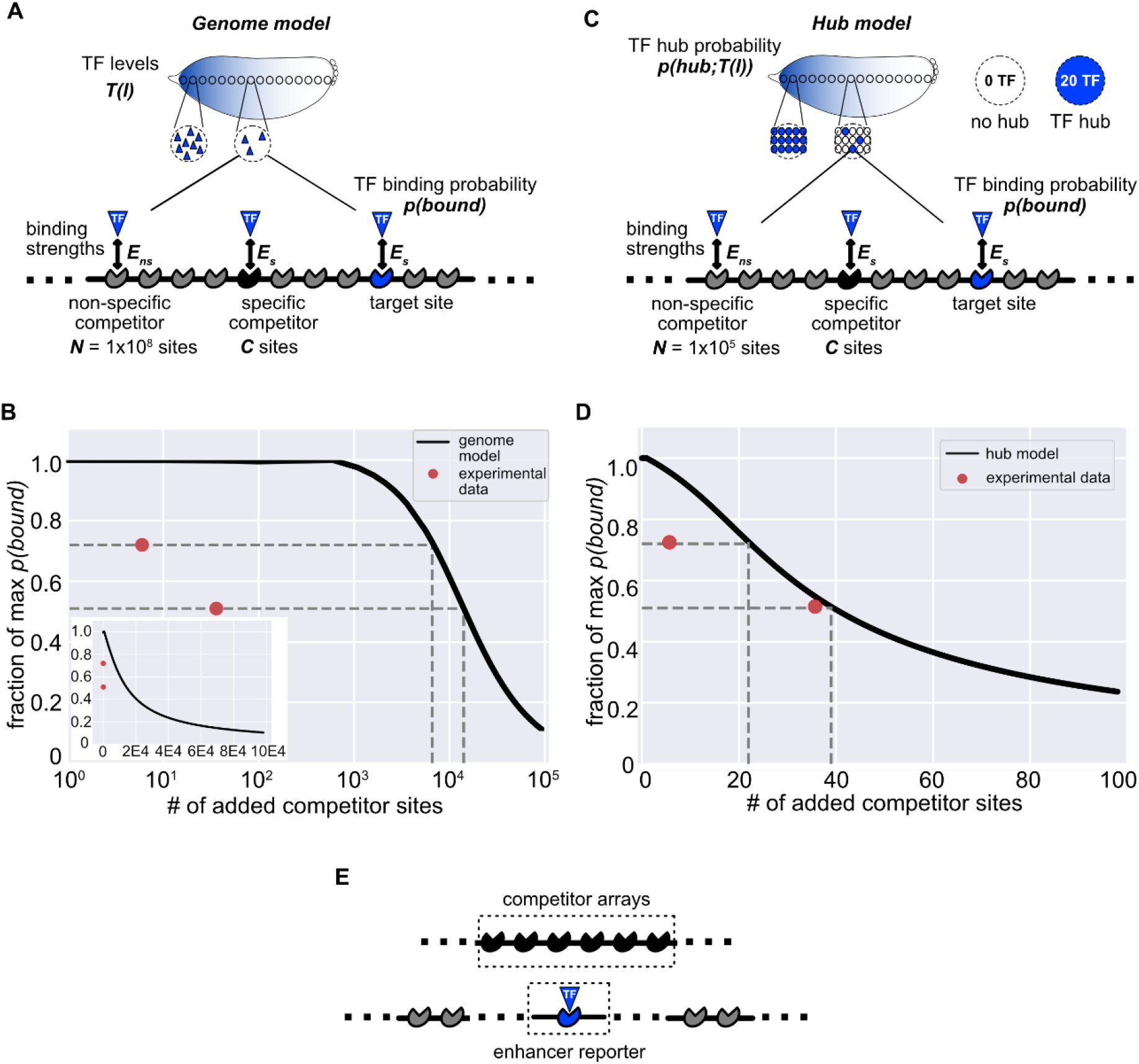
Modeling the impact of competitor binding sites on TF-enhancer binding. To understand how small transgenic sequences could induce the competition for TFs we observe, we created a thermodynamic model of TF binding at a single site as a function of TF levels, competitor sites, and binding strengths. **A**. Schematic of the parameters of the genome model where the whole genome is considered for TF binding. The probability of a TF molecule being bound at the target site, *p(bound)*, is determined by the parameters shown. The number of available TFs, *T*, varies as a function of embryo position *l* to match the measured Bcd gradient (Fowlkes, et al., 2008). We assume that all TF molecules are bound and can be bound to either the target site or competitor sites, which are divided into specific and non-specific sites. The number of non-specific sites, *N*, is held constant at 1×10^8^ while the number of specific competitor sites, *C*, is varied. TF molecules bind the target site and specific competitor sites with binding energy *Es* and bind non-specific sites with binding energy *Ens. Ens* is held constant at zero and *Es* is varied. With each set of parameters, *p(bound)* is calculated using equation 1 from the text. **B**. The fraction of maximum *p(bound)* as a function of number of added competitor sites using the genome model. *Es* is held constant at 10 and *l* is held constant at 27% embryo length. Model predictions are in black. Experimental data of the fraction of maximum hemizygous *hb*P2 reporter expression as a function of the number of Bcd binding sites in the transgene on the homologous chromosome is shown in red. Data points indicate the fraction of maximum *hb*P2 expression with a second *hb*P2 reporter, which contains 6 Bcd binding sites, or the 6xBcd array on the homologous chromosome, measured at 27% egg length. Dashed lines indicate the number of additional competitor sites predicted by the genome model to be required to produce the experimentally observed decrease in expression. The inset shows the same data on a linear x-axis. **C**. Schematic of the parameters of the hub model where TF binding is assumed to only occur within nuclear subregions. Each nucleus is divided into 1000 equally-sized regions, one of which contains the target site. As in the genome model, the output of the model is the probability of a TF molecule being bound at the target site, *p(bound)*. Based on previous measurements, the number of available TFs, *T*, is held constant at 20 for hub regions and 0 for non-hub regions (Mir, et al., 2017). Instead, the probability that a region in a nucleus is a hub is a function of embryo position *l* to match the Bcd gradient and we call this probability *p(hub; T(l))* (equation 2 in the text). As in the genome model, we assume all TFs are bound at the target site, competitor sites, or non-specific sites. In each region, the number of non-specific competitor sites, *N*, is 100,000 while the number of specific competitor sites, *C*, varies. The binding strength parameters *Es* and *E*_*ns*_ are the same as those used in the genome model. *p(bound)* is calculated as in the genome model using equation 1 from the text and multiplying the resulting value by *p(hub;T(l))*. This product is the final *p(bound)* value. **D**. Results of the hub model. The graph is as in B, with the fraction of maximum *p(bound)* as a function of the number of added competitor sites where the black line is the prediction of the hub model. As in B, *Es* is held at 10 and *l* is held constant at 27% egg length. Red points are the same experimental data as in B. Dashed lines indicate the number of additional competitor sites predicted by the hub model to be needed to produce the experimentally observed decrease in expression. **E**. Model results can be compared to experimentally measured decreases in reporter expression. The schematic shows how our models relate to our experimental system. The target site in the models is analogous to the enhancer-MS2 reporters in our experimental system. The added specific competitor sites of the models represent the TF binding site arrays or second reporter introduced on the homologous chromosome opposite the enhancer reporter. Although the exact relationship is not known, TF binding at enhancers is related to enhancer activity so the *p(bound)* output of our models is related to our measured enhancer reporter activity.

As we vary the parameters, we find that *p(bound; T(l))* changes in a qualitatively intuitive way. *p(bound; T(l))* decreases as a function of increasing competitor sites, decreasing difference in specific and non-specific binding strength, and decreasing TF levels (Supplemental Figure 6). To test the accuracy of our model, we compared our experimental measurements of expression changes as a function of additional Bcd binding sites to predicted changes in *p(bound; T(l))* as a function of added competitor sites. In our model, we assume there are 2000 specific competitor sites, based on experimental measurements of genome-wide Bcd binding in nc14 embryos (Hannon, et al., 2017), and then add additional competitor sites to mimic the addition of a reporter of Bcd binding site array. Our findings are similar even if we do not assume these “background” sites exist (Supplemental Figure 6). We recognize that the relationship between TF binding at an enhancer and gene transcription is complex (Grossman, et al., 2017; Chen, et al., 2020; Liu & Tijan, 2018) and do not expect our predicted *p(bound; T(l))* values to exactly predict gene expression levels. Still, gene expression is dependent on TF binding (Mir, et al., 2018; Shariati, et al., 2019) and so our *p(bound; T(l))* values provide useful ballpark estimates of how gene expression is expected to change as new competitor sites are introduced.

We compared our model predictions to the experimentally measured changes in activity of the *hb*P2 reporter. The *hb*P2 enhancer is a well-studied, Bcd-responsive enhancer and therefore makes a useful point of comparison for our model of Bcd binding (Driever & Nusslein-Volhard, 1989; Struhl, et al., 1989; Chen, et al., 2012). For simplicity, we compared our experimental data and model predictions at one position in the embryo, 27% egg length, where the *hb*P2 enhancer drives peak levels of expression in homozygous embryos. This means we hold *l*, and consequently *T*, constant and therefore refer to our model output as *p(bound)* from here on. To observe the effect specifically of introducing new specific competitor binding sites, we also used experimental measurements to estimate the difference between *Es* and *Ens* and held this constant (see Methods). In our experimental data, we see a 28% reduction in activity driven by the *hb*P2 reporter when a second reporter is present on the homologous chromosome. In contrast, our model predicts a 0.0003% decrease in *p(bound)* from the addition of 6 specific competitor sites, which is the number of known Bcd sites within the *hb*P2 reporter (Driever, et al., 1989). With this model, over 6,000 competitor sites are needed to get a 28% reduction in *p(bound)* (Figure 5C). Similarly, while we observe a 49% decrease in *hb*P2 expression in the presence of the 6xBcd array, which contains 36 Bcd binding sites, our model predicts only a 0.002% decrease in *p(bound)* with this number of added competitor sites. Based on model predictions, 14,100 specific competitor sites need to be added to achieve a 49% reduction in *p(bound)*. Thus, a simple thermodynamic model of molecular competition produces estimates a couple of orders of magnitude different from experimental measurements.

We suspected that the large discrepancy between our measured decreases in reporter activity and our model’s predictions of decrease in *p(bound)* are partially due to the model’s assumption that any Bcd molecule in the nucleus can bind the target site. Growing evidence indicates that TFs and other pieces of the transcriptional machinery are not distributed evenly throughout the nucleus, but instead tend to cluster in regions of high density, called hubs, separated by low density regions (Tsai, et al., 2019; Mir, et al., 2018; Cho, et al., 2018; Boehning, et al., 2018; Tsai, et al., 2017). This non-homogenous distribution seems functional, as transcription itself is also associated with these hubs (Tsai, et al., 2019; Chong, et al., 2018; Cho, et al., 2018). Compared to the whole nucleus, hubs have a higher concentration of TFs and a lower number of specific and non-specific binding sites. We predicted that the addition of a small number of binding sites, similar to the numbers found in our reporter constructs, may have a sizable impact on *p(bound)* in the context of individual TF hubs.

To test this, we modified our previous model (*genome model*) to look at the probability of TF binding at the same target site, assuming all TF binding happens within hubs (*hub model*). In our hub model, we divide the nucleus into 1000 hub-sized regions, based on the size of *Drosophila* embryonic nuclei and previous estimates of the distance between enhancers associated with the same TF hub (Tsai, et al., 2019; see Methods). Based on the measured distribution of distances between transcriptional spots in homozygous embryos, it is likely that reporters and TF binding site arrays transiently co-localize to the same hub-sized region (Supplemental Figure 7). Within each region, we assume there are 100,000 non-specific binding sites, which is the number of non-specific sites in the genome model (1×10^8^) divided by 1000.

The number of specific competitor sites is varied from 0 to 100. Based on previous measurements, the number of Bcd TFs present in a hub, *T*_*hub*_, is held constant at 20 molecules per hub, but the number of total Bcd molecules per nucleus, *T(l)*, follows the Bcd gradient along the embryo (Mir, et al., 2017). Regions that are not a hub are assumed to have 0 Bcd molecules.

Consequently, the *p(bound)* value in our hub model is found by multiplying the *p(bound)* value calculated using the same formula as the genome model (equation 1) by the probability that a given region is a TF hub (*p(hub; T(l));* equation 2). As in our genome model, we varied the difference between specific and non-specific binding energies.

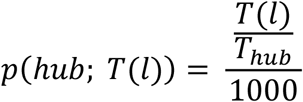

In comparison to the genome model, the hub model shows far better agreement with our experimental data. As with the genome model, we assume that some specific competitor sites already exist and ask how *p(bound)* changes as additional specific competitor sites are added. In the hub model, we assume the 2000 specific competitor sites of the genome model are evenly distributed throughout the genome and consequently two specific competitor sites are present in each sub-region of the nucleus. We again focus on one position in the embryo, 27% egg length, and therefore hold *T(l)* constant. Experimentally, we see a 28% decrease in the activity of the *hb*P2 reporter with the addition of a second *hb*P2 reporter. The hub model predicts a 5% decrease in *p(bound)* from the addition of the 6 Bcd binding sites in the *hb*P2 reporter (Figure 5D). Unlike the genome model, which requires over 6000 competitor sites for a 28% reduction in *p(bound)*, this magnitude reduction is achieved by 22 competitor sites in the hub model. With 36 competitor sites, the number of Bcd binding sites in the 6xBcd array, the hub model predicts a 46% decrease in *p(bound)* compared to the 49% decrease in *hb*P2 reporter expression we measure in the presence of the 6xBcd array.

It is notable the hub model better predicts the effect of a 6xBcd array than the second *hb*P2 reporter. While there are many simplifying assumptions in the model, for example, assuming *p(bound)* is proportional to expression output, there also is a key difference between the two experimental measurements. The 6xBcd array lacks a promoter, while the *hb*P2 reporter actively drives transcription. The model assumes that the only effect of adding the *hb*P2 reporter is the addition of competitor Bcd binding sites, but this reporter may also siphon away other key pieces of transcriptional machinery, which may explain why the measured effect of adding the *hb*P2 reporter is larger than predicted by either model.

## Discussion

Since the discovery of enhancers 40 years ago (Banerji, et al., 1981; Moreau, et al., 1981), transgenic reporters have been invaluable tools to study the principles governing *cis*-regulatory regions. With a few exceptions, it has largely been assumed that transgenic reporters do not meaningfully affect the expression of other genes. Here we challenge this assumption and investigate the observed competition between transgenic transcriptional reporters in developing *Drosophila* embryos. Using reporters controlled by different configurations of the *Kruppel* shadow enhancers, we show that expression of a single reporter is decreased in the presence of a second identical reporter. We further show that this effect is not limited to transgenic reporters, but that the expression of a nearby endogenous gene is also decreased in transgenic embryos.

Using non-transcribing arrays of TF binding sites, we find evidence that decreased reporter expression is due in part to decreased availability of key activating TFs of the *Kr* enhancers. Focusing on enhancer competition for the TF Bcd, we show that competitor Bcd binding arrays specifically affect the expression of Bcd-regulated enhancers, have a dosage-dependent effect on these enhancers, and shrink the width of the expression pattern of the enhancers. By developing a simple thermodynamic model, we predicted that the introduction of tens of additional Bcd binding sites can appreciably decrease gene expression, but only when TF binding is assumed to be limited to nuclear subregions.

### Transgenic reporters can affect the expression of other reporters and an endogenous gene

Due to the widespread use of transgenic reporters, we were surprised to find that our small reporters reduce the expression of not only other reporters, but a nearby endogenous gene. A deeper search of literature revealed that Laboulaye, et al., also found a distance-dependent decrease in endogenous gene expression in mice that is similar to our own results (Laboulaye, et al., 2018). While further systematic investigation is needed, these similar results in these distantly related organisms suggest that decreased endogenous gene expression may be a common consequence of transgenic reporters.

At first glance, our findings are also reminiscent of the transgene silencing previously reported in *Drosophila* (Kassis, et al., 1991; Kassis, 1994; Pal-Bhadra, et al., 2002). Pioneering studies found that flies containing multiple copies of a transgene showed reduced expression of the transgene as well as the corresponding endogenous gene (Pal-Bhadra, et al., 1997; Pal-Bhadra, et al., 1999). This silencing was shown to depend on Polycomb-mediated repression in the case of transgenes, and post-transcriptional RNAi mechanisms in the case of the endogenous gene (Pal-Bhadra, et al., 2002). While our findings share some similarities with transgene silencing, and may well rely on related mechanisms, numerous differences lead us to believe we are observing a distinct phenomenon. Unlike these previous studies, our transgenic flies, which contain a mini-*white* marker, do not show lighter eye color in homozygotes compared to hemizygotes (Supplemental Figure 8). This suggests that overall expression from our transgenic insertion sites is not being ubiquitously repressed. If our observations were only the consequence of silencing mechanisms, triggered by increased amounts of transgenic DNA, we would expect to see larger reporter competition effects with our duplicated enhancer constructs compared to the single enhancer versions. Instead, for both the distal and proximal enhancers, we see a trend of larger decreases in homozygous expression levels compared to hemizygous expression levels with the single enhancer constructs (Figure 1B). Further, unlike the findings of Pal-Bhadra, our transgenic reporter decreases expression of an endogenous gene with which it does not share sequence homology (Pal-Bhadra, et al., 2002).

### Reporters and non-transcribed DNA sequences can induce competition for TFs

In addition to the studies described above, which describe how a transgene can alter the expression of genes, there are several studies that describe how the presence of non-transcribing pieces of DNA can alter expression. Work in flies and mouse cells showed that highly repetitive genomic sequences can alter gene expression, likely by binding and sequestering TF molecules away from their target genes (Liu, et al., 2007; Janssen, et al., 2000). In yeast, repetitive sequences of “decoy” tetO TF binding sites can change the relationship between tetO levels and the expression of a gene regulated by tetO (Lee & Maheshri, 2012), and individual competitor binding sites in bacteria also have a similar effect (Brewster, et al., 2014). These studies underscore the regulatory importance of repetitive non-coding DNA sequences, which make up the majority of many genomes, by titrating available TF levels.

The repetitive sequences investigated in these studies are much longer (6Mb of major satellite DNA in mice, 7Mb of satellite V DNA in flies) than our transgenic constructs, which all contain less than 5kb of regulatory sequence and 10s of TF binding sites (Guenatri, et al., 2004; Janssen, et al., 2000b). The large effect our transgenic constructs have on gene expression levels are therefore initially surprising. It is easier to imagine how very large, repetitive DNA sequences could sequester meaningful amounts of TFs than small sequences containing as few as six TF binding sites. In particular, this competition is surprising because we observe it even in the peak regions of the reporter expression patterns, where we expect activating TF levels to be high. For example, there are 250,000 molecules of the TF Zld per nucleus in the embryo (Biggin, 2011; BNID 106849, Milo et al., 2010), yet we see evidence of competition by introducing only six new strong Zld binding sites.

We suspect that the effect of our reporters and binding site arrays on expression levels, as well as the effect of large repetitive sequences, partially stems from the non-uniform distribution of TFs in the nucleus. Although heterogeneity in the nucleus has been long observed with DNA (Nagele, et al., 1999; Manuelidis, 1984), recent studies revealed that TFs and other pieces of the transcriptional machinery are also distributed unevenly throughout the nucleus (Tsai, et al., 2019; Mir, et al., 2018; Cho, et al., 2018; Boehning, et al., 2018; Tsai, et al., 2017). There are several potential consequences of the organization of TFs into hubs. First, if our competitive binding arrays end up outside of a so-called TF “hub” with the enhancer reporter (or nearby endogenous genes), TF levels functionally available to the enhancer may be low enough to disrupt reporter activity. Second, if our binding arrays and reporters are found in the same hub, they may be competing for a fairly small pool of TFs. Previous measurements suggest there are roughly 20 Bcd molecules per hub (Mir, et al., 2017). Lastly, the presence of a binding array may affect the properties of the hubs themselves. Another study showed that the deletion of TF-recruiting enhancers can decrease TF hub size and therefore lower gene expression (Tsai, et al., 2019). In addition, Zld plays a key role in the formation of Bcd hubs (Mir, et al, 2018), suggesting that our Zld binding site arrays may sequester both Zld and Bcd molecules.

Several aspects of our data support the hypothesis of local competition. First, reporters that spend more time in close physical proximity in the nucleus compete more than reporters that are further apart (Supplemental Figure 5). Similarly, the endogenous gene *Piezo*, whose expression is decreased in the presence of the duplicated distal reporter, is within the same topologically associating domain (TAD) as the insertion site of the transgene during nc14 (Hug, et al., 2017; Supplemental Figure 9). This suggests that *Piezo* and the reporter likely inhabit the same nuclear subregion and have access to the same local pool of TFs.

### Thermodynamic model of TF binding implicates TF hubs in competition

In line with our experimental data, our modeling results suggest that local competition for TFs is consistent with the observed decrease in expression levels. To rationalize how the addition of a small number of competitor TF binding sites could meaningfully decrease expression levels, we developed two simple thermodynamic models of TF binding. Our hub model, which assumes all TF binding is restricted to nuclear subregions matches our experimental data more closely than the genome model, which assumes that all TF molecules have access to the whole genome. Our findings suggest an unexplored consequence of TF hubs. Previous studies have shown that TF hubs help to increase local concentration of TFs to increase gene expression (Tsai, et al., 2019; Chong, et al., 2018). Here, we show the flip side of this coin -- the non-uniform distribution of TFs can also induce competition among binding sites. We note that we have used only strong binding sites in our competitor arrays and plan to test the effect of binding arrays consisting of non-optimal TF binding sites, as enhancers containing sub-optimal binding sites have been shown to be important for establishing TF hubs (Tsai, et al., 2019). There may be a balance between sequestering TFs, as we see here, and recruiting TFs to a local region that could depend on the affinity of the binding sites present.

Our goal in developing a simple model of our system was to generate ballpark predictions about the behavior of the system, using experimentally measured parameters and minimal assumptions. While our hub model better matches our experimental findings than the genome model, we recognize that it is a simplification of reality and as such cannot fully describe our system. For example, we assume that existing specific competitor sites are evenly distributed throughout the genome, but in reality, chromatin accessibility and the clustering of TF binding sites in cis-regulatory regions alters the distribution of available binding sites (Berman, et al., 2002; Li, et al., 2011). The “true” number of specific competitor sites in any given sub-nuclear region will vary and consequently *p(bound)* of a given target site will depend on the surrounding sequences in the same region (Supplemental Figure 10). With these simplifying assumptions, come an incomplete ability to explain some experimental data. We find that expression levels driven by the duplicated distal reporter significantly decrease in the presence of the smallest and largest of our Bcd binding site arrays, but, unexpectedly, are not affected by the presence of the intermediate sized array (Supplemental Figure 3). We do not fully understand this observation, but suspect that it has to do with the exact recruitment of TFs and other molecular factors mediated by this combination of DNA sequences.

### Implications for transgenic reporters and underlying biology

Our work adds to the evidence that transgenic reporters can have measurable effects on endogenous gene expression (Laboulaye, et al., 2018; Pal-Bahdra, et al., 1997) and also builds on our understanding of the mechanisms behind this phenomenon. We note that our transgenic fly lines develop without any gross phenotypic defects in ideal laboratory conditions, making it tempting to assume that any effects of transgenic reporters are negligible. While much about the mechanisms and effects of reporters on endogenous gene expression remains to be discovered, our findings provide some practical lessons for using transgenic reporters. First, investigators should use caution in interpreting changes in expression levels or patterns when comparing assays using one reporter to those using multiple reporters simultaneously. We find clear evidence that our reporters compete with one another when present in the same nucleus and as this seems to be mediated by competition for TFs, we suspect this finding is true beyond our specific reporters and system. Additionally, potential effects of reporters on nearby endogenous gene expression should be considered in study design and data interpretation.

Beyond the implications for the use of transgenic reporters, our findings suggest that the distribution of TF binding sites, both in the genome and in 3D space, is a potential tuning mechanism for dose-response relationships between TF levels and target genes. Previous studies in bacteria and yeast have shown that competitor TF binding sites can modulate the doseresponse relationship of TF levels and gene expression, and that this modulation depends on the relative affinity of competitor versus gene-regulating TF binding sites (Brewster, et al., 2014; Lee & Maheshri, 2012). This effect may generate an unappreciated selection pressure to either retain or eliminate TF binding sites that are not directly regulating a specific target gene. The observations of TF sequestration across a wide range of organisms suggest that this phenomenon is conserved and likely plays a functional role in regulating gene expression.

## Acknowledgements

We thank Hernan Garcia for flies containing MCP-GFP and His-RFP transgenes, as well as for useful discussion of our observations. We thank Lily Li for helpful suggestions on the model and all members of the Wunderlich lab for feedback on the project. We also thank German Enciso, Anthony Long, Thomas Schilling, and Rahul Warrior for helpful feedback and suggestions on the project.

## Author Contributions

**RW:** conceptualization, software, formal analysis, investigation, writing - original draft, writing - review & editing, visualization. **MG:** Investigation, resources, writing - review & editing. **ZW:** conceptualization, resources, writing - review & editing, supervision, funding acquisition.

## Methods

### Generation of transgenic fly lines

Transgenic fly lines containing an enhancer-MS2 reporter were generated by phiC31-mediated insertion into the second or third chromosome, as described in Waymack, et al., 2020. Unless otherwise indicated, all reporter constructs and TF binding site arrays were integrated into the same site on the second chromosome via phiC31-mediated integration. These constructs were injected into y[1] w[1118]; PBac{y[+]-attP-3B}VK00002 (BDRC stock #9723) embryos by BestGene Inc (Chino Hills, CA). For the reporter constructs inserted into chromosome 3, plasmids were injected into y[1] w[1118]; PBac{y[+]-attP-3B}VK00033 (BDRC stock #9750) embryos by BestGene Inc (Chino Hills, CA).. The *Kruppel* enhancer reporters contained a single, duplicated, or shadow enhancer pair and the *Kruppel* promoter upstream of 24 MS2 repeats and a *yellow* reporter gene cloned into the pBphi vector (Garcia, et al., 2013). These are the same enhancer-MS2 reporters as used and described in Waymack, et al., 2020. The *hunchback* P2 enhancer reporter is that used in Garcia, et al., 2013 and consists of the *hunchback* P2 enhancer and P2 promoter upstream of 24 MS2 repeats and a *lacZ* reporter (Garcia, et al., 2013). Exact genomic sequences used in each reporter construct are given in Supplementary file 1.

Hemizygous embryos were generated by crossing male flies homozygous for an enhancer-MS2 reporter to females expressing RFP-tagged histones and MCP-GFP (Garcia, et al., 2013).

Homozygous embryos were generated by crossing virgin females of the F1 hemizygous offspring just described with males homozygous for the same enhancer-MS2 reporter. Embryos with one copy of an enhancer reporter and one copy of a TF binding site array were generated by crossing the virgin female hemizygous offspring (i.e. containing one enhancer-MS2 reporter allele) with males homozygous for the corresponding TF binding site array.

### Generation of TF binding site arrays

To generate our competitor binding site arrays for the four different TFs investigated, we started with the sequence of the *hb* P2 enhancer, which is well known to be Bcd responsive and contains six Bcd binding sites (Driever & Nusslein-Volhard, 1989; Struhl, et al., 1989; Chen, et al., 2012). This 236bp sequence was our 1xBcd array and the starting point for our other binding site arrays. To generate the Hb, Zld, and Stat92E arrays used we modified the six Bcd binding sites of the *hb* P2 enhancer to be the consensus motif for the corresponding TF (while retaining the same 236bp total length of the array) and had these sequences synthesized by Integrated DNA Technologies Inc (San Diego, CA). Previously defined consensus motifs were used for Zld (Xu, et al., 2014), Hb (Stanojevic, et al., 1989), and Stat92E (Yan, et al., 1996). To generate the 3x and 6xBcd arrays we performed Golden Gate assembly to ligate three or six copies of the 1xBcd array together with 10bp random sequences between each repeat, to avoid potential repeat removal during transformations. All of the described TF binding site arrays were inserted into the same plasmid backbone, which was a modified version of the pBphi vector used for our enhancer-MS2 reporters, which lacks any enhancers, promoters, or MS2 sequence. We generated this vector by removing the *Kr* distal enhancer, *Kr* promoter, MS2 cassette, *yellow* sequence, and termination sequence from our distal MS2 reporter through digestion with NotI and XbaI. We then used Gibson assembly to ligate in the appropriate TF binding site array to this backbone. Sequences for the TF binding site arrays are provided in Supplemental File 1.

### Embryo preparation and image acquisition

Living embryos were collected and dechorinated before being mounted onto a permeable membrane in halocarbon 27 oil and placed under a glass coverslip as in Garcia, et al., 2013. Individual embryos were then imaged as described in Waymack, et al., on a Nikon A1R point scanning confocal microscope using a 60X/1.4 N.A. oil immersion objective and laser settings of 40uW for 488nm and 35uW for 561nm (Waymack, et al., 2020). To track transcription, 21 slice Z-stacks, at 0.5um steps, were taken throughout the length of nc14 at roughly 30s intervals. To identify the imaged position in the embryo, the whole embryo was imaged after nc14 prior to gastrulation at 20X using the same laser power settings. This whole embryo image was used to assign each transcription spot into one of 42 bins across the anterior-posterior (AP) axis of the embryo. The first bin corresponds to the anterior end of the embryo.

### Measurement of transcriptional reporter activity

Tracking of nuclei and MCP-GFP bound MS2 transcriptional spots was done using the image analysis Matlab pipeline described in Garcia et al., 2013, which can be accessed at the Garcia lab Github (https://github.com/GarciaLab/mRNADynamics). Calling of transcriptional bursts to use for analysis was done as in Waymack et al., 2020. In short, transcriptional traces captured during nc14 consisting of at least three points were used for analysis. To measure total mRNA produced by all of our reporter configurations, we integrated the area under the curve of the transcriptional spot’s fluorescence across the time of nc14 (Supplemental Figure 1).

For every tracked spot of transcription, background fluorescence at each time point is estimated as the offset of fitting the 2D maximum projection of the Z-stack image centered around the transcriptional spot to a gaussian curve, using Matlab *lsqnonlin*. This background estimate is subtracted from the raw spot fluorescence intensity. The resulting fluorescence traces across the time of nc14 are then subject to smoothing by the LOWESS method with a span of 10%. The smoothed traces were used to measure transcriptional parameters and noise. Traces consisting of fewer than three time frames were removed from calculations. The area under each smoothed transcriptional trace is integrated using the Matlab *trapz* function, which gives the total integrated fluorescence value for that transcriptional spot. This integrated fluorescence is proportional to the number of transcripts produced by an enhancer reporter (Garcia, et al., 2013; Lammers, et al., 2020). We group all transcriptional spots of a given reporter configuration by AP bin (position in the embryo) and calculate the average total integrated fluorescence value in each AP bin. For each reporter configuration we identify the AP bin with the highest average integrated fluorescence value as the region of peak expression. In the text, unless otherwise indicated, the integrated fluorescence or peak expression values correspond to the average integrated fluorescence value at this AP bin (Supplemental Figure 1).

### qRT-PCR to measure expression of endogenous genes in varying genetic backgrounds

Flies were allowed to lay eggs on molasses plates for 2.5 hours, so that most embryos collected were in nc14. Flies were either homozygous for the duplicated distal reporter on chromosome 2 or of the same genetic background (BDSC #9723) but did not have the transgene. The embryos collected from each plate were pooled and total RNA was extracted and purified using TRIzol (Thermo Fisher Scientific) and the Direct-zol RNA Miniprep kit (Zymo Research). cDNA was generated using SuperScript III RT Kit (Thermo Fisher Scientific). qPCR amplification was then done using the TaqMan Gene Expression Master Mix (Thermo Fisher Scientific). The data for each group, transgenic or non-transgenic, is from three separate biological replicates (i.e. the colored circles in Figure 4C are biological replicates), each done in technical triplicates. Relative RNA levels of each measured gene was calculated using the 2^-ddC(t)^ method, using *RpII140* as the reference gene. The TaqMan FAM probes used for each gene were DM01803576_g1 for *Piezo*, Dm01825813_g1 for *Mcr*, Dm01803642_g1 for *Btk29A*, and DM02134593_g1 for *RpII140*

### Description of the genome model of TF binding

We developed a simple thermodynamic model that looks at the probability of a TF molecule being bound at a single target site. For simplicity, we assume all TF molecules are bound either specifically or non-specifically. The probability of TF being bound at the target site, *p(bound)*, is then determined by the number of TF molecules, *T(l)*, the number of non-specific competitor binding sites, *N*, the number of specific competitor sites, *C*, and the difference in specific versus non-specific binding energies, *E*_*s*_ and *E*_*ns*_ respectively. The number of TF molecules, *T(l)*, follows the Bcd gradient (Fowlkes, et al., 2008) and is determined by position in the embryo, *l*, with a maximum value of 20,000 at the anterior tip of the modeled embryo (Biggin, 2011). For ease of comparison with our experimental data, we only consider binding probability at one position (*l* = 27% egg length) and thereby hold *T* constant, unless otherwise indicated. We hold the number of non-specific binding sites, *N*, constant at 1×10^8^. Using statistical mechanics, we first enumerated the possible states of our system and their associated Boltzman weights (Bintu, et al., 2005). In these states, x indicates the number of TFs that are bound at specific competitor sites.

| TF binding configuration | State | Statistical weight |
| --- | --- | --- |
| Non-specific binding only | $\frac{N^T}{T!}$ | $e^{-TE_{ns}}$ |
| Competitor sites and non-specific binding | $\sum_{x=0}^C \frac{N^{T-x}}{(T-x)!} \times \frac{C!}{x!(C-x)!}$ | $e^{-[(T-x)E_{ns}+xE_s]}$ |
| Target site, competitor sites and non-specific binding | $\sum_{x=0}^C \frac{N^{T-1-x}}{(T-1-x)!} \times \frac{C!}{x!(C-x)!}$ | $e^{-[(T-1-x)E_{ns}+xE_s+E_s]}$ |

With all of the possible states of the system and the associated statistical weights, we can calculate *p(bound)* by dividing the statistical weight of the state with TF bound at the target site by the combined statistical weights of all possible states:

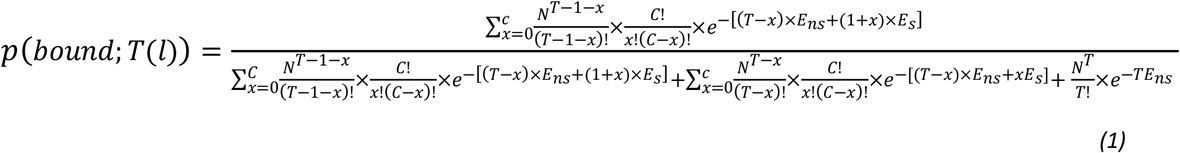

With this equation we can calculate the probability of a TF molecule being bound at the target site for a given number of TF molecules, specific competitor sites, and difference in binding affinity at specific vs non-specific sites. In the main text (Figure 5), we assume 2,000 background competitor sites already exist and span a range of 0 to 100,000 added specific competitor sites. To facilitate comparison with our experimental data, we looked at binding probability at one position in the embryo by holding *l* and consequently *T* constant. We focus on the effect of adding specific competitor binding sites and as such hold the difference between *E*_*ns*_ and *E*_*s*_, and *T(l)* constant. *E*_*ns*_ is held at 0kBT and *E*_*s*_ is held at -10kBT, based on previously measured differences in specific vs non-specific DNA binding (Jung, et al., 2018; Bintu, et al., 2005). Specifically, the formula 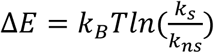 was applied to the measured binding affinities, from Jung, et al., of TFs to their consensus sequence and highly mutated consensus sequences (Jung, et al., 2018). *T(l)* is held at 5468, corresponding to 27% egg length in the modeled embryo. In Supplemental Figure 6, we explore how *p(bound)* changes as a function of our other parameters (i.e. *T(l)*, the difference between *E*_*ns*_ and *E*_*s*_, or the number of background specific competitor sites).

### Description of the hub model of TF binding

Calculation of *p(bound)* in the hub model is similar to the genome model but assumes all TF molecules are restricted to “hub” regions (sub-regions of the nucleus containing a high concentration of TFs) and therefore takes into account the probability that the nuclear sub-region containing the target site is a TF hub. In the hub model, each nucleus is divided into 1000 equal-sized regions with a radius of 256nm. This estimate for the size of nuclear regions was based on the average distance between interacting loci found by Tsai, et al., (Tsai, et al., 2019) of approximately 360nm, the approximate volume of *Drosophila* embryonic nuclei of 70um^3^, and an estimation of the amount of DNA contained within a TF hub of the size seen in Tsai, et al. The nuclear volume of 70um3 was reached by estimating the nucleus to be a sphere and using the formula 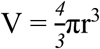 with r = 2.5um (estimated from imaging data).

As the nucleus is divided into 1000 hub regions, we set the number of non-specific binding sites, N, to 100,000, which is 1/1000th of the value in the genome model (10^8^). For simplicity, we assume that DNA is distributed uniformly in the nucleus and as such the amount of DNA in each region is the same, which also allows us to have the same number of total non-specific binding sites per nucleus as in the genome model (10^5^ x 1,000 hubs = 10^8^). To maintain the same number of total specific competitor sites as the genome model, we assume 2 background specific competitor sites per region and vary the number of added specific competitor sites per region from 0 to 100. These values were reached by dividing the number of background or added specific competitor sites from the genome model by 1,000 (2,000 / 1,000 = 2 and 100,000 / 1,000 = 100). Based on the observation of Mir, et al., (Mir et al., 2017) that the number of Bcd molecules per hub did not change along the Bcd gradient, we hold *T* constant at 20 if a region is a TF hub or 0 if it is not a TF hub. To account for this additional condition of whether the target site is within a TF hub or not, we calculate the probability that the region containing the target site is a TF hub:

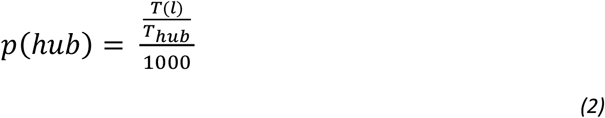

Where *T*_*hub*_ is 20 TF molecules found in a hub and *T(l)* is the total number of TFs in the nucleus, as determined by the Bcd gradient (Fowlkes, et al., 2008). To obtain the final *p(bound)* value from the hub model the *p(hub)* value of equation 2 is multiplied by the value obtained using the above parameters in equation 1.

### Plotting p(bound) as a function of added specific competitor sites

To best simulate our experimental system, where we add transgenic constructs containing Bcd binding sites into a genome that already contains Bcd binding sites, we focused on how *p(bound)* changes as a function of added specific competitor sites. We therefore hold *l*, and consequently nuclear TF levels *T*, constant. Similarly, we hold constant the difference in binding energy between specific and non-specific sites by setting *E*_*s*_ to -10 and *E*_*ns*_ to 0. To account for specific Bcd binding sites that already exist in the *Drosophila* genome, we set the *p(bound)* value when a set number of “background” competitor sites exist as our reference maximum *p(bound)* value.

We estimated the total number of true Bcd binding sites in nc14 embryos to be 2,000 based on the number of genome-wide Bcd ChIP-seq peaks reported by Hannon, et al (Hannon, et al., 2017). Therefore for the genome model, the graph shown in Figure 5B depicts how *p(bound)* changes as a function of additional specific competitor sites beyond 2,000. For the hub model, we assume that these 2,000 Bcd binding sites are equally distributed throughout the genome and consequently there are two specific Bcd binding sites per nuclear region (2,000 / 1,000 = 2). The graph in Figure 5D shows how *p(bound)* changes as a function of additional specific competitor sites per nuclear region beyond the baseline two.

### Statistical Methods

To determine statistical differences in levels of competition (Figures 1 and 4) and expression boundaries (Figure 3 and Supplemental Figure 3) between reporters we performed bootstrapping to estimate 95% confidence intervals. To do so, we randomly sampled with replacement the integrated fluorescence values of all of the transcriptional spots tracked in the AP bin of peak expression for both the hemizygous and homozygous configurations of the respective enhancer reporters. We averaged this value for the hemizygous configuration and for the homozygous configuration and then divided this average homozygous expression by the hemizygous expression to get our competition value (i.e. % hemizygous expression). This was done 1,000 times and each time the difference between the competition value found using the original real data set and that found using the randomly resampled data was calculated. We then took the top and bottom 2.5 percentiles of these differences as our upper and lower error bounds, respectively.

We estimated the error in expression boundaries in a similar fashion. We again perform 1,000 rounds of bootstrapping by randomly sampling, with replacement, the integrated fluorescence values from rows of transcriptional spots along the AP embryo axis for a given enhancer construct. Each column corresponds to a single AP bin in the embryo. We randomly sampled rows equal to the total number of rows in the original data set and using these found the anterior-most and posterior-most AP bins that produce greater than or equal to 50% of the maximum expression measured in the hemizygous configuration of that reporter. Empirical 95% confidence intervals were calculated as above by finding the 2.5th and 97.5th percentiles of the distribution of differences between the expression boundaries found using the original data and those found using each iteration of resampled data.

To determine whether the distal enhancer reporter produces significantly lower expression levels in the presence of a non-transcribing distal enhancer (Figure 1C) we performed a t-test comparing the integrated fluorescence values recorded in the region of peak expression of the two configurations.

## Supplemental Materials

Supplemental File 1 - Fasta file containing the sequences of all enhancer sequences used in reporters as well as the sequences of the TF binding site arrays.

Matlab and Python code used for analysis and generation of figures can be found at github.com/WunderlichLab/TFCompetitionCode

---

**Supplemental Figure 1 –.**
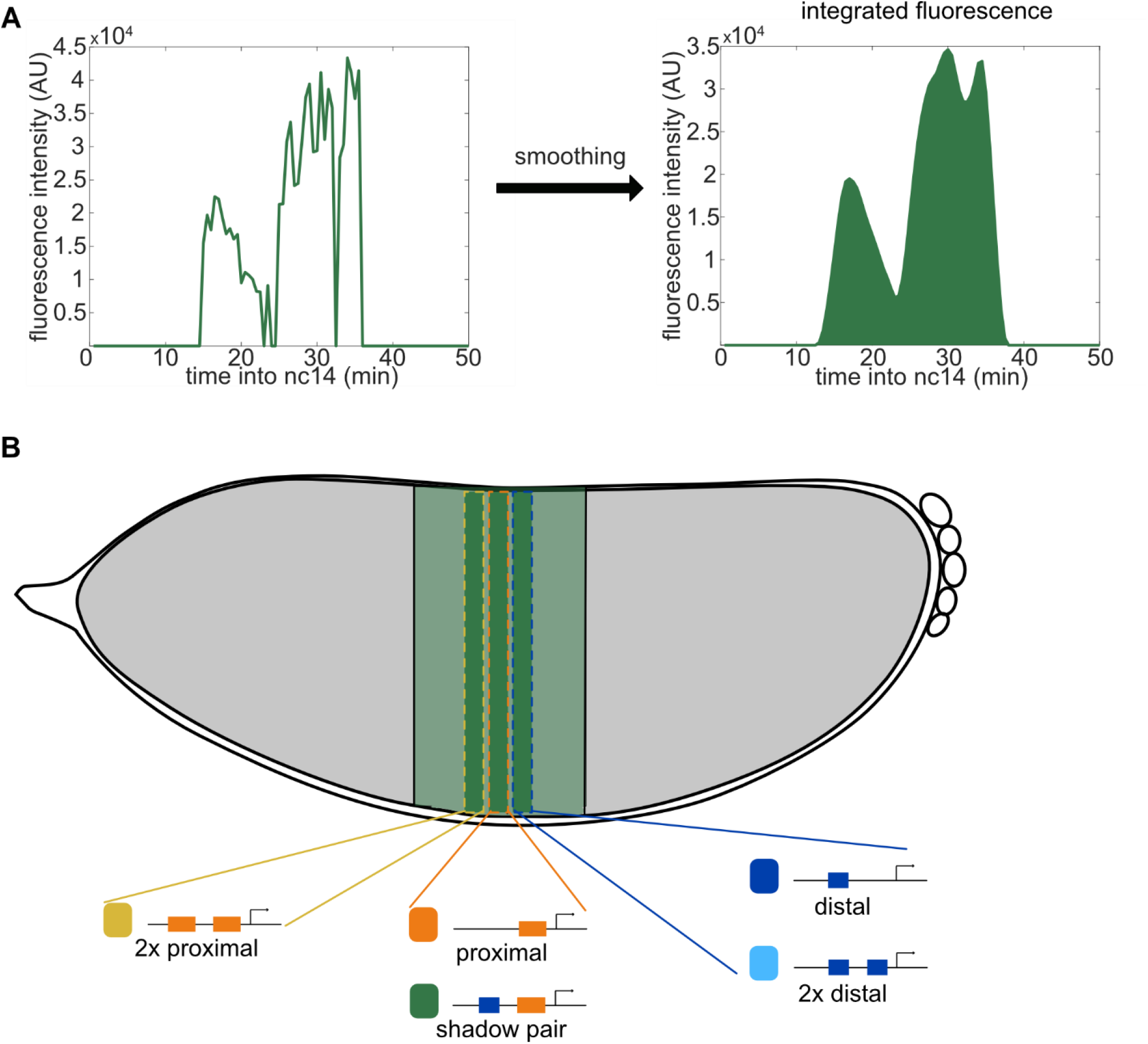
**Total enhancer activity is measured by integrating the area under fluorescence time trace curves at the region of peak expression**. To measure the activity of our enhancer reporters, we track individual spots of transcription across the time of nc14. Our enhancer reporters drive expression of 24 repeats of MS2 sequence that when transcribed forms stem loops that are bound by the fluorescently tagged coat protein, MCP-GFP. This enables us to visualize and track sites of nascent transcripts as spots of fluorescence above background (Garcia, et al., 2013). **A**. Transcriptional traces are first smoothed using the Lowess method (see Methods for more details). Integrated fluorescence, which is proportional to total mRNA produced and hence indicative of enhancer activity, is measured by calculating the area under the fluorescence curve. This area is calculated with the trapz function in Matlab using the fluorescence points recorded during the first 50 minutes of nc14. **B**. Total expression for each enhancer reporter is calculated at the position of that reporter’s peak expression levels along the anterior-posterior axis of the embryo. We divide the embryo along the anterior-posterior axis into 41 equally sized bins that each encompass 2.5% of the embryo and calculate the average total reporter expression per bin. Schematic shows the *Kr* expression pattern during nc14 as the green stripe at the center of the embryo. Within this stripe, the different enhancer reporters have slightly different regions of peak expression as indicated by the darker green rectangles. The enhancer reporter or reporters that drive peak expression in each region is indicated below the corresponding bin. The duplicated proximal reporter shows peak expression at 47.5% egg length (0% corresponding to the anterior tip of the embryo), the proximal and shadow pair reporters at 50% egg length, and the single and duplicated distal reporters at 52.5% egg length.

**Supplemental Figure 2 –.**
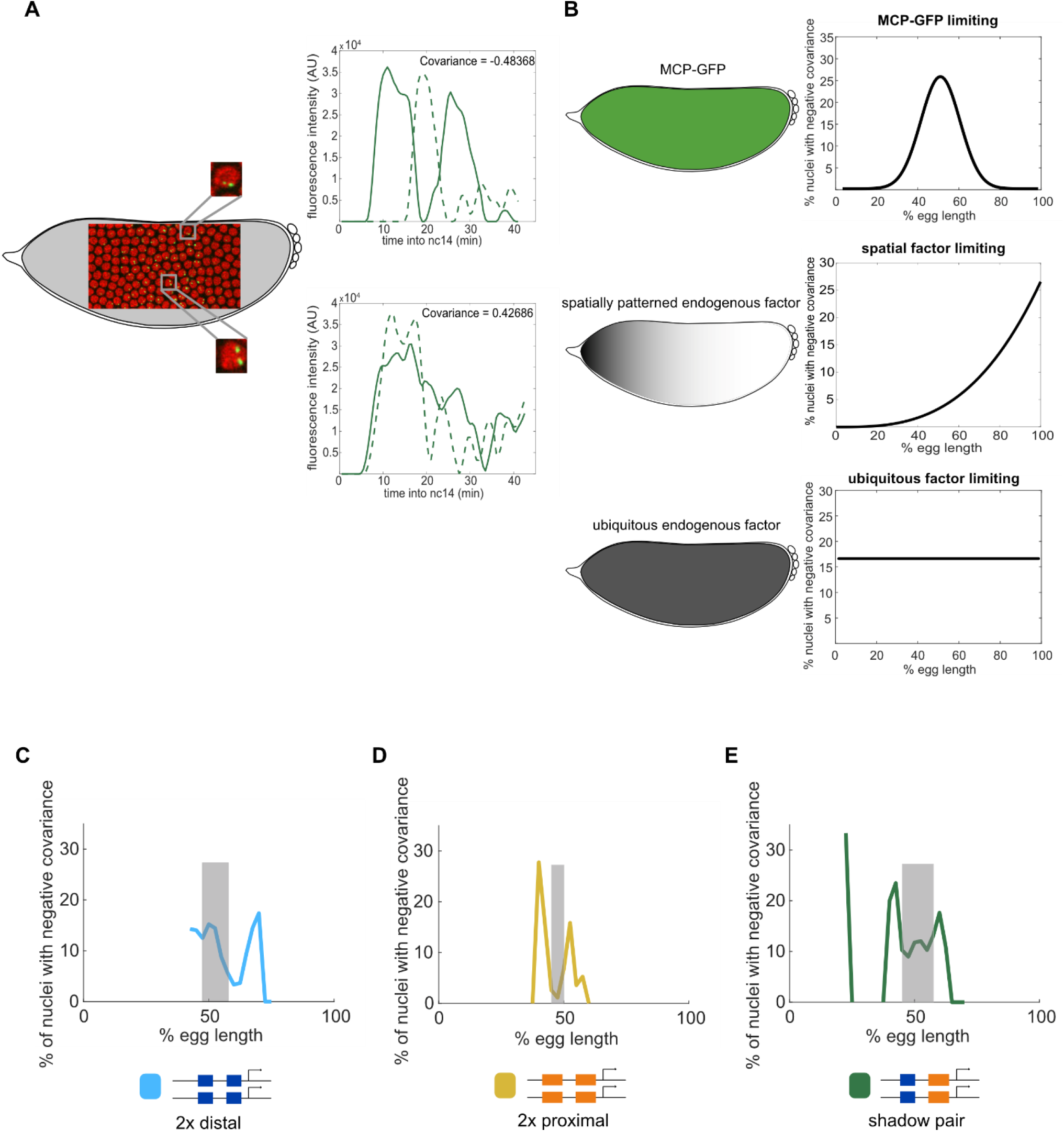
Patterns of negative covariance suggest competition for spatially-patterned factor. To assess whether the differences in reporter expression levels observed between homozygous and hemizygous embryos stem from competition for factors required for reporter visualization, we measured the covariance of the activity of identical reporters in individual nuclei. A fraction of nuclei display negative covariance, which is indicative of an antagonistic relationship between the activities of the two reporters. **A**. Still image from a live imaging movie where nuclei are colored in red and sites of active transcription are green spots. Insets show zoomed-in example nuclei that display negative covariance (top) or do not (bottom). The graphs to the right show the transcriptional activity of the two reporters in each nucleus across the time of nc14. **B**. Schematics of expression patterns of different possible limiting factors and the corresponding expected patterns of negative covariance. The MCP-GFP reporter is expressed ubiquitously across the length of the embryo, while an endogenous factor may be expressed ubiquitously or in a spatial pattern. Graphs to the right show expected spatial pattern of negative covariance rates if reporters are competing for limiting levels of MCP-GFP (top), a spatially patterned endogenous factor (middle), or a ubiquitously expressed endogenous factor (bottom). **C-E**. The fraction of nuclei that display negative covariance as a function of egg length. Grey highlight indicates the region of 75% max expression for that reporter construct. **C**. Duplicated distal. **D**. Duplicated proximal. **E**. Shadow pair.

**Supplemental Figure 3 –.**
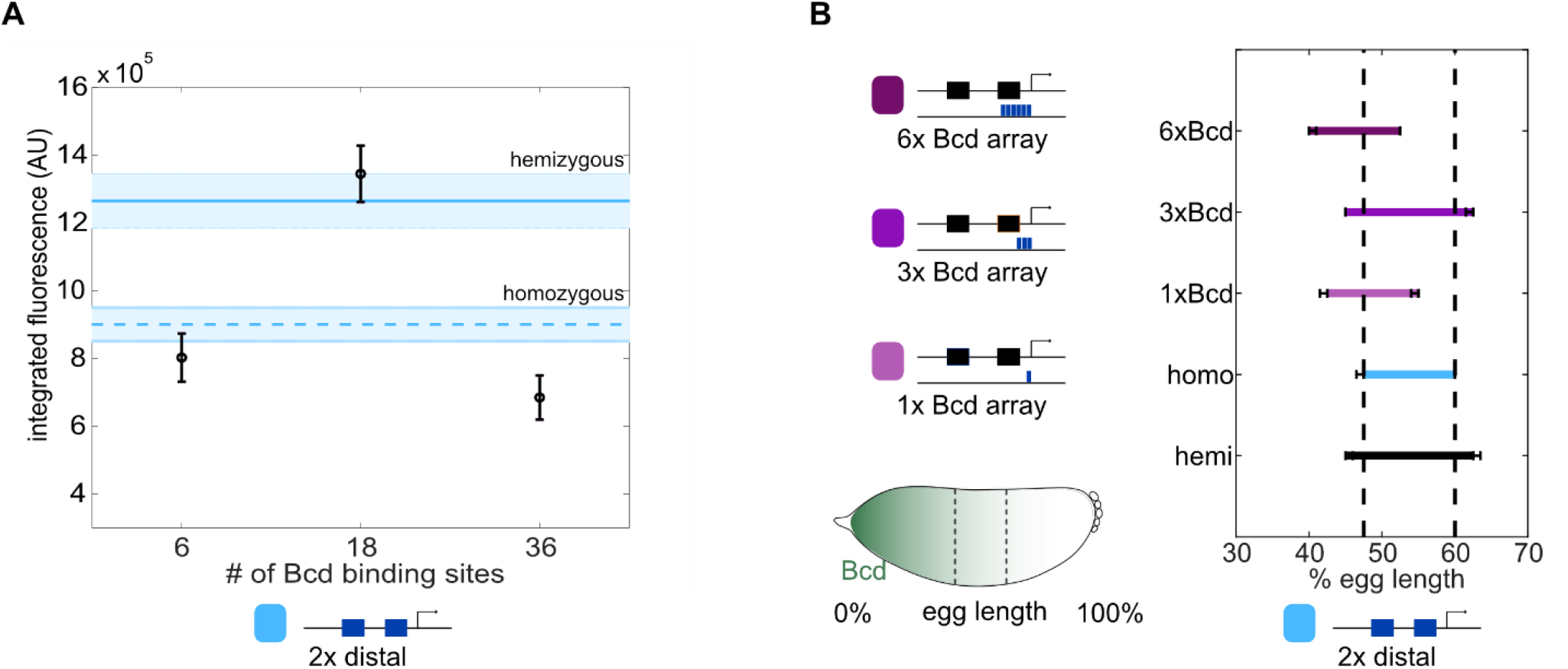
Competitor Bicoid binding sites decrease and shift the activity of the duplicated distal reporter. To assess the effect of increasing number of competitor Bcd sites on the activity driven by the duplicated distal enhancer construct, we measured its expression in the presence of larger Bcd binding site arrays (the same as used in Figure 3). **A**. Duplicated distal reporter expression changes non-linearly with increasing number of Bcd binding sites. Graph shows the total expression driven by the duplicated distal reporter at its region of peak expression in the presence of Bcd binding arrays with the indicated number of Bcd binding sites. Horizontal lines indicate peak expression levels per allele in hemizygous (top) and homozygous (bottom) embryos. Error bars and shading indicate 95% confidence intervals. In the presence of the 1xBcd array, peak expression of the duplicated distal reporter is decreased 37% relative to hemizygous levels and is further reduced 46% relative to hemizygous levels in the presence of the 6xBcd array. Interestingly, with the 3xBcd array the duplicated distal reporter’s activity is not significantly affected, which we do not fully understand but propose may be due to the molecular composition of the microenvironment created around the duplicated distal reporter with the 3xBcd array. **B**. The expression domain of the duplicated distal reporter shifts in the presence of competitor Bcd binding arrays and these shifts qualitatively match changes in peak expression levels. The graph shows the range of the expression pattern of the indicated construct to 50% of peak expression levels of the homozygous configuration, whose boundaries are indicated with dashed vertical lines. Error bars represent 95% confidence intervals from 1000 rounds of bootstrapping.

**Supplemental Figure 4 –.**
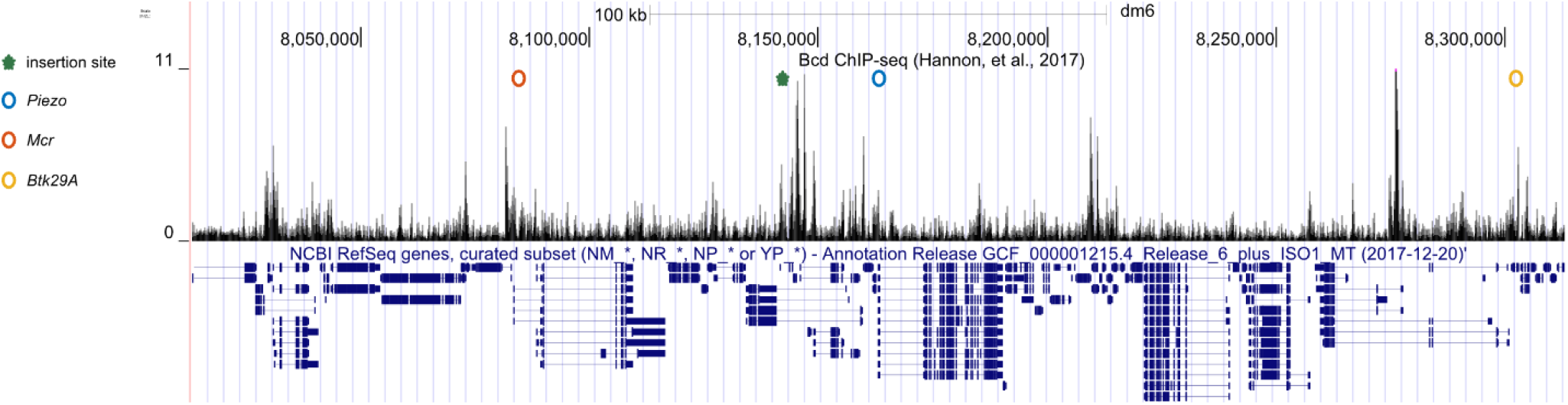
Bcd binding surrounding genes near chromosome 2 transgenic insertion site. To assess the potential effect of our enhancer-MS2 transgenes on endogenous gene expression, we measured the expression of *Piezo, Mcr*, and *Btk29A* in embryos with or without the duplicated distal transgene inserted on chromosome 2. We looked at these three genes due to their proximity to the transgenic integration site (Figure 4) and the observation that they are all likely regulated by Bcd. All three genes have expression patterns in the early embryo that are consistent with activation by Bcd and previous Chip-seq in nc14 embryos indicates Bcd binding surrounding these genes (Hannon, et al., 2017). Figure shows UCSC genome browser window of 300kb centered on the *Piezo* transcription start site. Top line shows genomic coordinates, black peaks indicate Bcd binding as measured in Hannon, et al., 2017, and bottom in blue indicates gene annotations. Green star marks site of chromosome 2 attP insertion site. Circles mark the transcription start sites of the three endogenous genes where *Piezo* is blue, *Mcr* is red, and *Btk29A* is yellow.

**Supplemental Figure 5 –.**
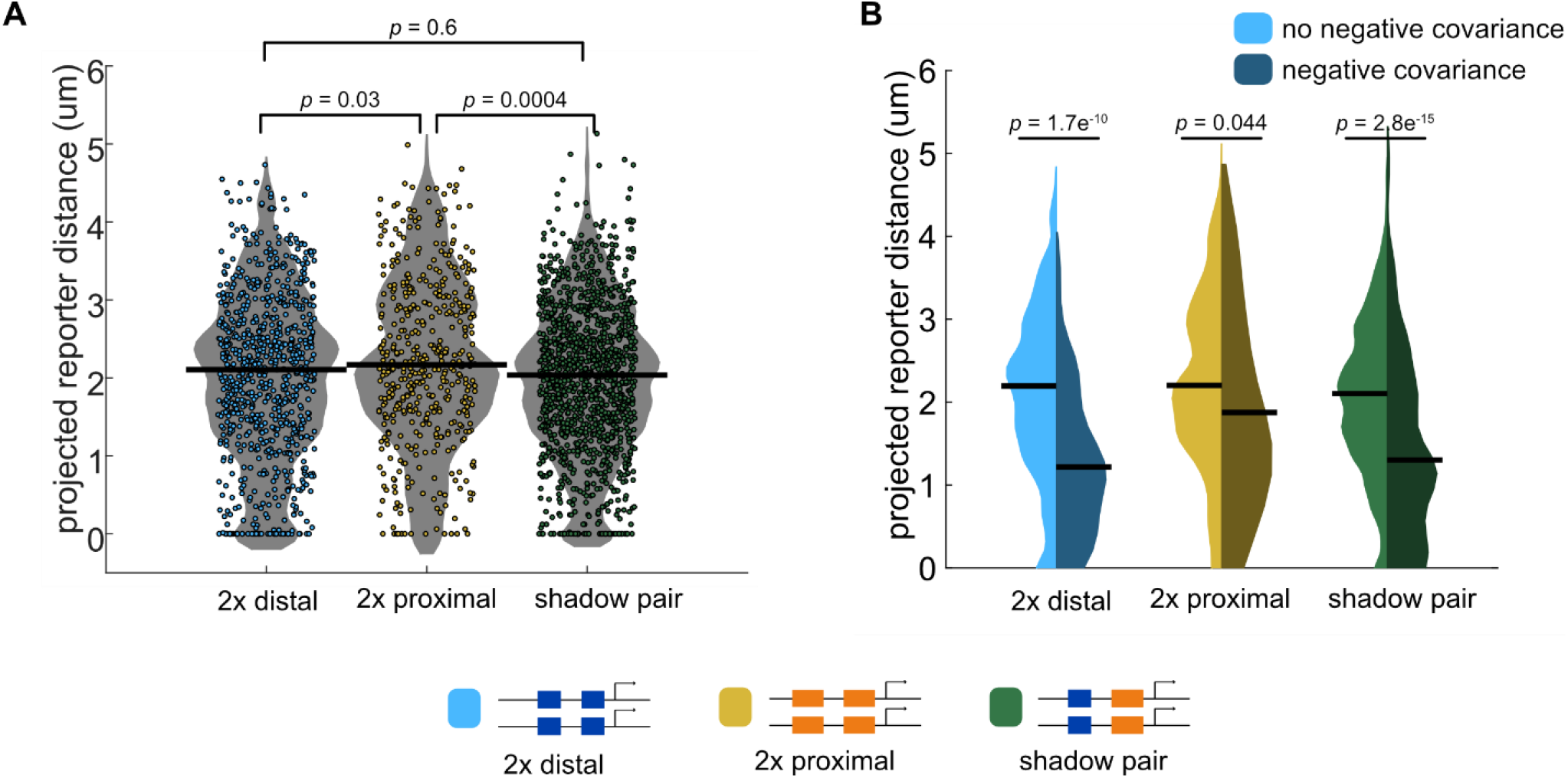
Average distance between reporters is negatively correlated with competition. To test the hypothesis that reporters are competing for locally limited TFs, we compared the average distance between our reporters in a nucleus with the degree of competition that reporter shows. **A**. Graph shows the average projected distance between identical reporters in the same nucleus for the indicated reporters. Each colored circle represents the average distance between the two reporters in a single nucleus across the time of nc14. The duplicated distal and shadow pair reporters, which show significant competition (Figure 1A) on average are much closer together in the nucleus than are the duplicated proximal reporters, which do not show significant competition. Horizontal lines indicate medians. Significance determined by Kruskal-Wallis test with Bonferroni correction. **B**. Graph shows the distributions of projected distances between identical reporters in a nucleus broken down into nuclei with or without negative covariance, which is indicative of reporter competition. The left, lighter colored half of each violin plot shows the distribution of average reporter distance during nc14 in nuclei that do not show negative covariance. Right half of each violin plot shows this distribution in nuclei with negative covariance. Horizontal lines indicate medians. In all three reporter constructs, nuclei whose reporters display negative covariance, which is indicative of competition, have reporters that are on average significantly closer together than are reporters that do not display negative covariance. Significance determined by Kolmogorov-Smirnov test.

**Supplemental Figure 6 –.**
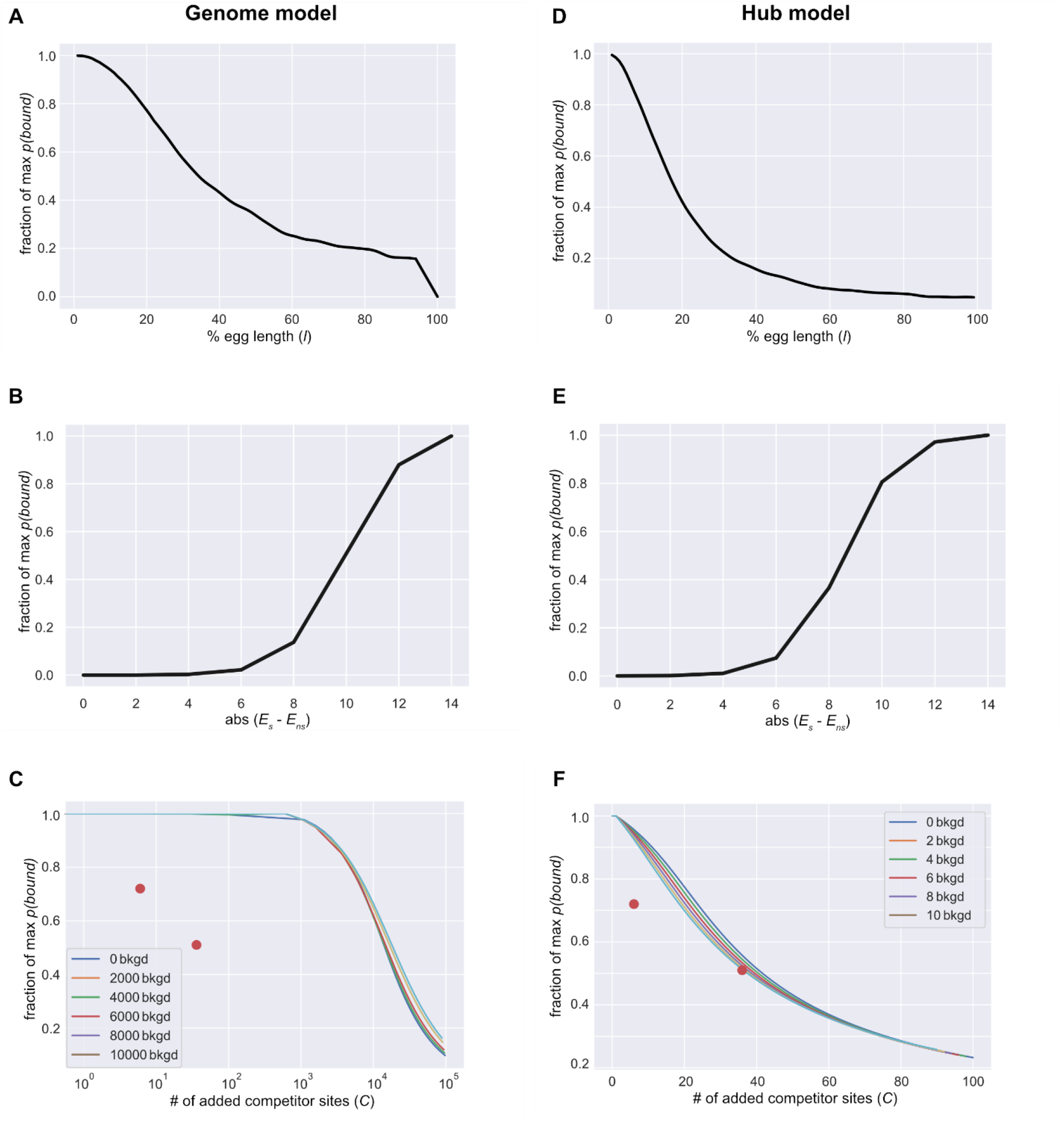
How *p(bound)* changes with varying parameters. To get a sense for how our model behaves, we looked at how *p(bound)* changes as a function of our three parameters (TF levels, binding energy difference, and number of specific competitor sites) for both our genome and hub models. **A**. In the genome model, *p(bound*) decreases with decreasing levels of TF. Graph shows model prediction of *p(bound)* as a function of embryo position, *l*, which determines TF levels following the Bcd gradient. We set the maximum Bcd level to 20,000 at the anterior of the embryo (Biggin, 2011) and use this value with the measured Bcd gradient (Fowlkes, et al., 2008) to estimate TF levels at each point along the embryo. 0% egg length corresponds to the anterior of the embryo. Number of specific competitor sites is held constant at 2,000 and the difference in specific vs non-specific binding energies is held constant at 10. **B**. In the genome model, binding probability increases as the difference between specific and non-specific binding energies increases. Graph shows genome model prediction of the fraction of maximum binding probability as a function of the difference between specific and non-specific binding energies. Non-specific binding energy is held constant at 0 while specific binding energy is decreased. Here the number of TFs is held constant at 5,468 (corresponding to *l* = 27% egg length) and the number of specific competitor sites is held constant at 2,000. **C**. The sensitivity of binding probability to added competitor sites in the genome model is dependent on the number of pre-existing or “baseline” specific competitor sites. Graph shows the fraction of maximum binding probability as a function of added specific competitor sites. The different colored lines show model predictions depending on the number of baseline specific competitor sites. Red points show experimental data of the fraction of peak hemizygous expression for the *hb*P2 reporter as homozygotes (top point) or with the 6xBcd array (bottom point). The number of TFs is held constant at 5,468 (corresponding to *l* = 27% egg length) and the difference in specific versus non-specific binding energies is held constant at 10. **D**. Binding probability predicted by the hub model decreases along the length of the embryo. The graph shows the fraction of maximum binding probability of the hub model as a function of embryo position, which in the hub model determines the probability of a nuclear subregion being a TF hub. The number of specific competitor sites is held constant at 2 and the difference in binding energies between specific and non-specific sites is held constant at 10. **E**. Graph as in B, showing the predictions of the hub model. Like with the genome model, binding probability increases with increasing difference between specific and non-specific binding energies. Non-specific binding energy is held constant at 0 while specific binding energy is decreased. Here embryo position *l*, and consequently *p(hub;T(l)*), is held constant at 27% egg length and the number of specific competitor sites per region is held constant at 2. **G**. As with the genome model, the sensitivity of binding probability to added competitor sites in the hub model is dependent on the number of pre-existing specific competitor sites. Graph is as in C for predictions of the hub model depending on the indicated number of baseline specific competitor sites. Red points are the same experimental data as in C. Here, embryo position *l*, and consequently *p(hub;T(l)*), is held constant at 27% egg and the difference in binding energies between specific and non-specific competitor sites is held constant at 10.

**Supplemental Figure 7 –.**
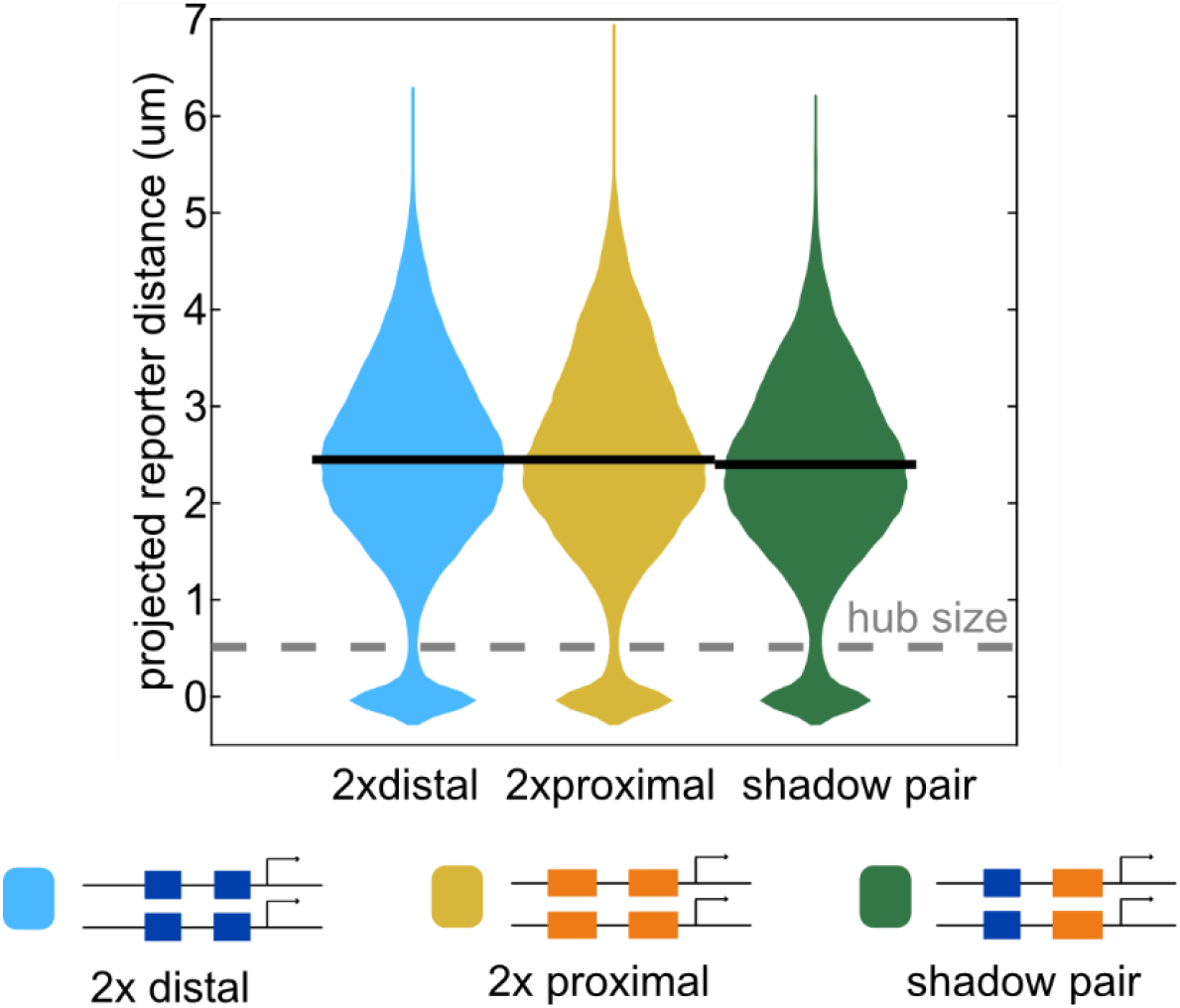
Distribution of all measured distances between transcriptional reporters. To assess whether enhancer reporters were close enough to be accessing the same “hub” region within the nucleus, we looked at the distribution of reporter distances in homozygous embryos of the three indicated reporter constructs. Violin plots show the distribution of projected distances between transcriptional spots for every time point in nc14 where two transcriptional spots were tracked in a nucleus, across all measured nuclei in each of the indicated constructs. Gray dashed line indicates the 512nm diameter of nuclear regions in our hub model. For all three constructs, the two copies of the reporter are within the same hub-sized region in between 7% and 8% of all recorded time points, based on estimates of TF hub size (Tsai, et al., 2019; see Methods for more details).

**Supplemental Figure 8 –.**
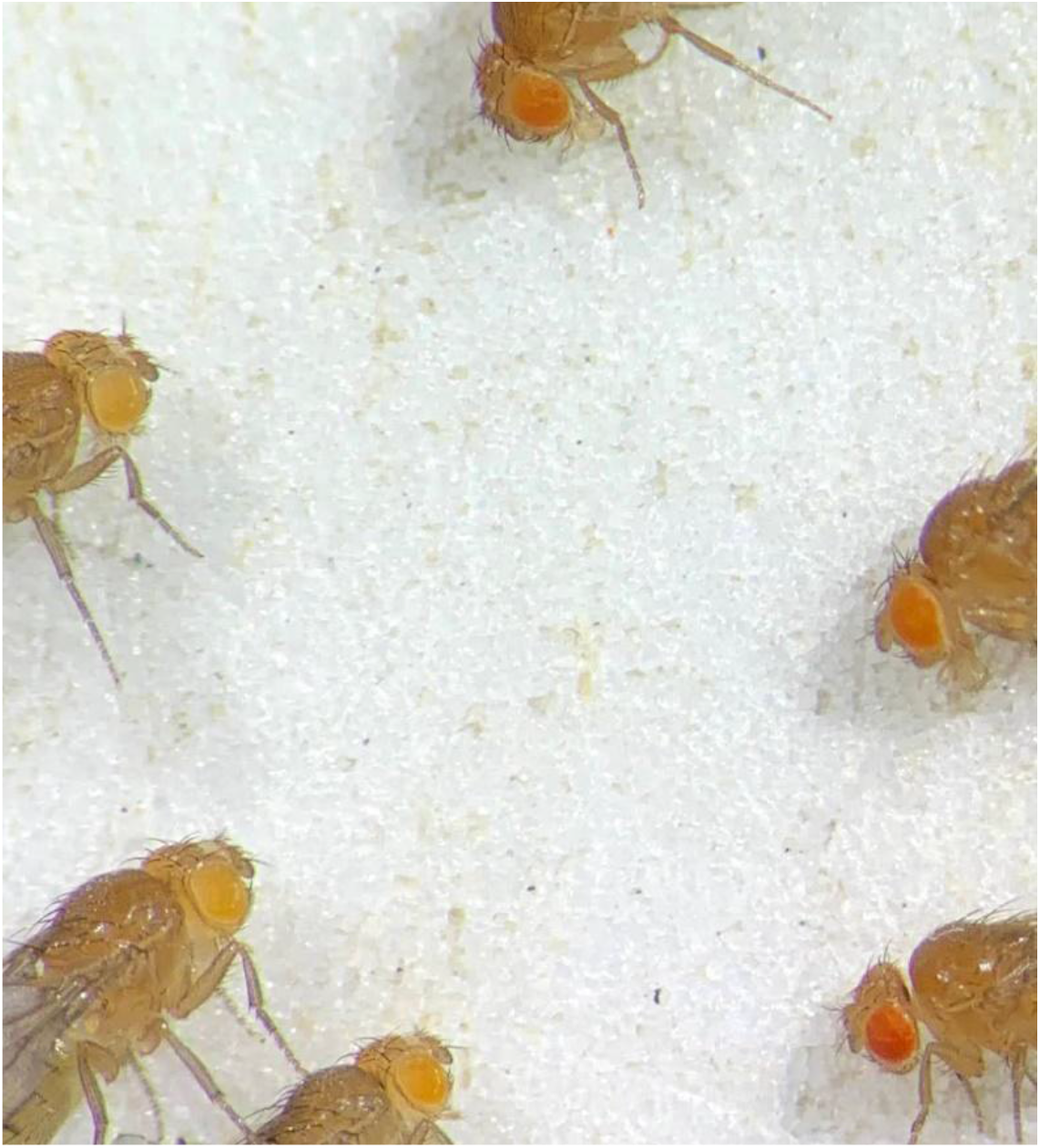
Comparison of homozygous and hemizygous fly eye color. Unlike previous observations of transgene silencing (Pal-Bhadra, et al., 1997), flies homozygous for our enhancer reporters, which contain a mini-*white* marker, do not show lighter eye color than hemizygous flies. The three flies on the left are hemizygous for the shadow pair reporter on chromosome 2 and the three flies on the right are homozygous for this same reporter. On both sides, the top two flies are female and the bottom-most fly is male.

**Supplemental Figure 9 –.**
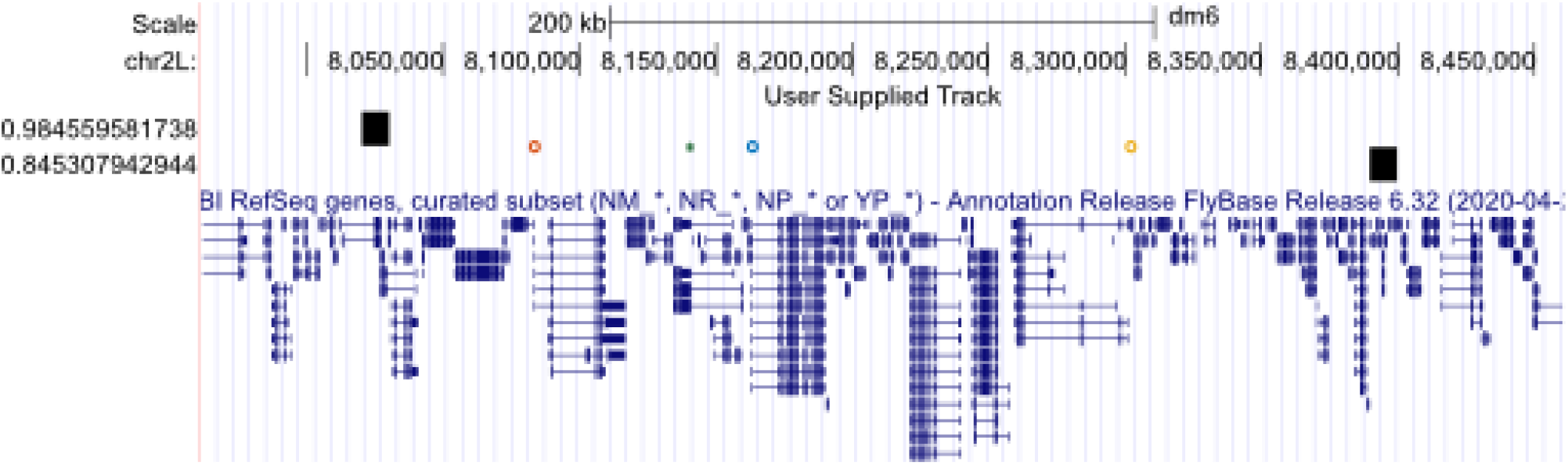
Transgenic insertion site and affected endogenous gene fall within the same TAD. Ours and previous data suggest that transgenes have a larger effect on the expression of endogenous genes closer to the transgene on a linear piece of DNA (Laboulaye, et al., 2018). As the genome is organized three dimensionally, we suspect that the distance between transgenes and endogenous genes in 3D space is important in determining the effect of a transgenic reporter on endogenous gene expression. As such, we used previously published Hi-C data from nc14 embryos to ask whether our transgene is likely contained within the same TAD as the measured endogenous genes. The figure shows a UCSC genome browser window with our transgenic insertion site marked as a green star and the three measured endogenous genes, *Piezo, Mcr*, and *Btk29A*, indicated as the blue, red, and yellow circles, respectively. The closest TAD boundaries as measured in Hug, et al., 2017 are indicated as black rectangles (Hug, et al., 2017). The transgenic insertion site and all three endogenous genes are contained within the same TAD during nc14.

**Supplemental Figure 10 –.**
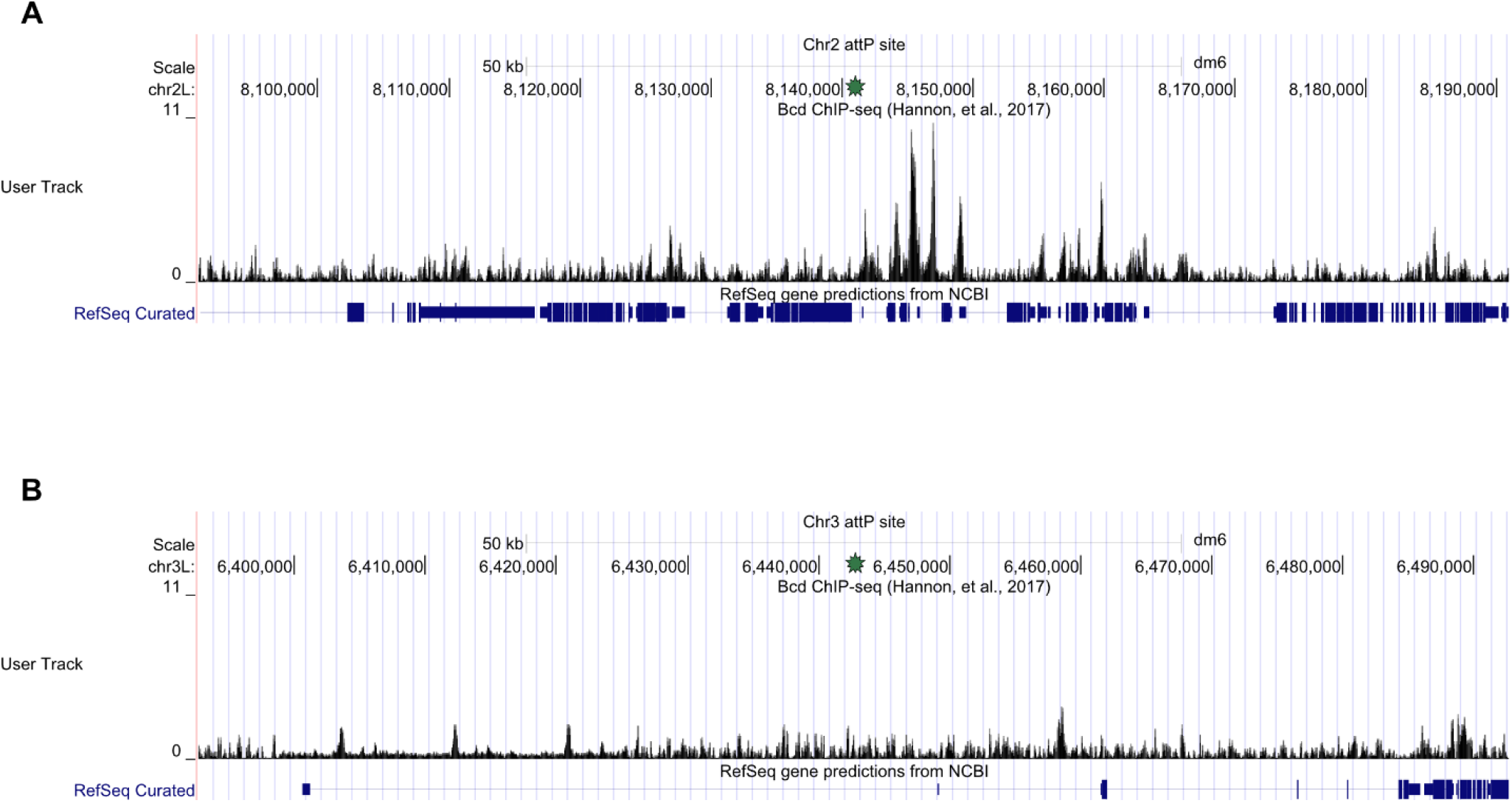
Bcd binding surrounding the two transgenic insertion sites. Our hub model predicts that TF competition is in part dependent on the number of existing TF binding sites within the same nuclear subregion. In our hub model we assume that all existing TF binding sites are distributed uniformly throughout the genome, but in reality, this is not the case. This can be seen with the Bcd binding surrounding our two transgenic insertion sites. **A**. UCSC genome browser window of 100kb centered on the attP insertion site used on chromosome 2. Top line shows genomic coordinates and black peaks indicate Bcd ChIP-seq reads from Hannon, et al., 2017 (Hannon, et al., 2017). **B**. UCSC genome browser as in A, but centered on the attP insertion site used on chromosome 3. Green star in A and B indicates the genomic coordinates of the respective insertion sites.

